# MrvR, a Group B *Streptococcus* Transcription Factor that Controls Multiple Virulence Traits

**DOI:** 10.1101/2020.11.17.386367

**Authors:** Allison N. Dammann, Anna B. Chamby, Andrew J. Catomeris, Kyle M. Davidson, Hervé Tettelin, Jan-Peter van Pijkeren, Kathyayini P. Gopalakrishna, Mary F. Keith, Jordan L. Elder, Adam J. Ratner, Thomas A. Hooven

## Abstract

*Streptococcus agalactiae* (group B *Streptococcus*; GBS) remains a dominant cause of serious neonatal infections. One aspect of GBS that renders it particularly virulent during the perinatal period is its ability to invade the chorioamniotic membranes and persist in amniotic fluid, which is nutritionally deplete and rich in fetal immunologic factors such as antimicrobial peptides. We used next-generation sequencing of transposon-genome junctions (Tn-seq) to identify five GBS genes that promote survival in the presence of human amniotic fluid. We confirmed our Tn-seq findings using a novel CRISPR inhibition (CRISPRi) gene expression knockdown system. This analysis showed that one gene, which encodes a GntR-class transcription factor that we named MrvR, conferred a significant fitness benefit to GBS in amniotic fluid. We generated an isogenic targeted knockout of the *mrvR* gene, which we found to have a growth defect in amniotic fluid relative to the wild type parent strain. In addition to growing poorly in amniotic fluid, the knockout also showed a significant biofilm defect *in vitro*. Subsequent *in vivo* studies showed that, while the knockout was able to cause persistent murine vaginal colonization, pregnant mice colonized with the knockout strain did not develop preterm labor despite consistent GBS invasion of the uterus and the fetoplacental units. In contrast, pregnant mice colonized with wild type GBS consistently deliver prematurely. Similarly, in a sepsis model in which 87% of mice infected with wild type GBS died within three days, none of the mice infected with the knockout strain died. In order to better understand the mechanism by which this newly identified transcription factor controls GBS virulence, we performed electrophoresis mobility shift assays with recombinant MrvR and whole-genome transcriptomic analysis on the knockout and wild type strains. We show that MrvR binds to its own promoter region, suggesting likely self-regulation. RNA-seq revealed that the transcription factor affects expression of a wide range of genes across the GBS chromosome. Nucleotide biosynthesis and salvage pathways were highly represented among the set of differentially expressed genes, suggesting a linkage between purine or pyrimidine availability and activity of MrvR in multiple GBS virulence traits.

## Introduction

*Streptococcus agalactiae* (group B *Streptococcus*; GBS) is a cause of chorioamnionitis, stillbirth, and neonatal infections including bacteremia, pneumonia, and meningitis (1–10). GBS is a common commensal of the intestinal and reproductive tracts in healthy adults, among whom invasive disease is rare (11, 12). In the pregnant or neonatal host, however, GBS can be highly invasive, breaching anatomic and immunologic barriers with potentially severe consequences (13, 14).

GBS is known to express a number of virulence factors such as adhesins (15–17), IgA binding proteins (18–20), and the cytotoxic ornithine-rhamnopolyene *β*-hemolysin/cytolysin (13, 21–30). Many of these are regulated by transcription factors, some of which have been studied and described in detail (31–40), yet many of the predicted transcription factors encoded by GBS remain minimally characterized.

We have previously described development of a GBS saturated transposon mutant library and its use in Tn-seq experiments to identify essential and conditionally essential genes (41, 42). In this study, we performed Tn-seq on GBS grown in human amniotic fluid in order to identify bacterial genes that promote survival in the nutrient-poor, unhospitable growth conditions of the amniotic sac.

Among the candidate genes was a transcription factor with domain features marking it as a member of the GntR protein superfamily. We named this GntR-class transcription factor MrvR (multiple regulator of virulence). Here we describe the role of MrvR in several GBS virulence-related phenotypes, including survival in human amniotic fluid, *in vitro* biofilm formation, and invasive bacteremia and preterm labor in murine models. We also use transcriptomic data and electrophoresis mobility shift assay results to demonstrate autoregulation of MrvR expression and its significant role in modifying expression of genes integral to nucleotide metabolism.

## Results

### Five GBS genes are conditionally essential for GBS survival in human amniotic fluid

Using a previously described transposon mutant library in an A909 (serotype Ia) background (41), we performed Tn-seq after growth challenge in eight human amniotic fluid samples from amniocentesis (two to three technical replicates in three separate biological samples). Prior to use for Tn-seq, the amniotic fluid was filter sterilized to remove any contaminating microorganisms and host cells that may have been present. The same mutant library was grown in rich media as an input control.

We used ESSENTIALS, a publicly available Tn-seq bioinformatics package, to analyze sequencing data from the Tn-seq experiment. This analysis revealed five candidate conditionally essential genes (**Figure 1 and Table 1**).

**Figure 1.**
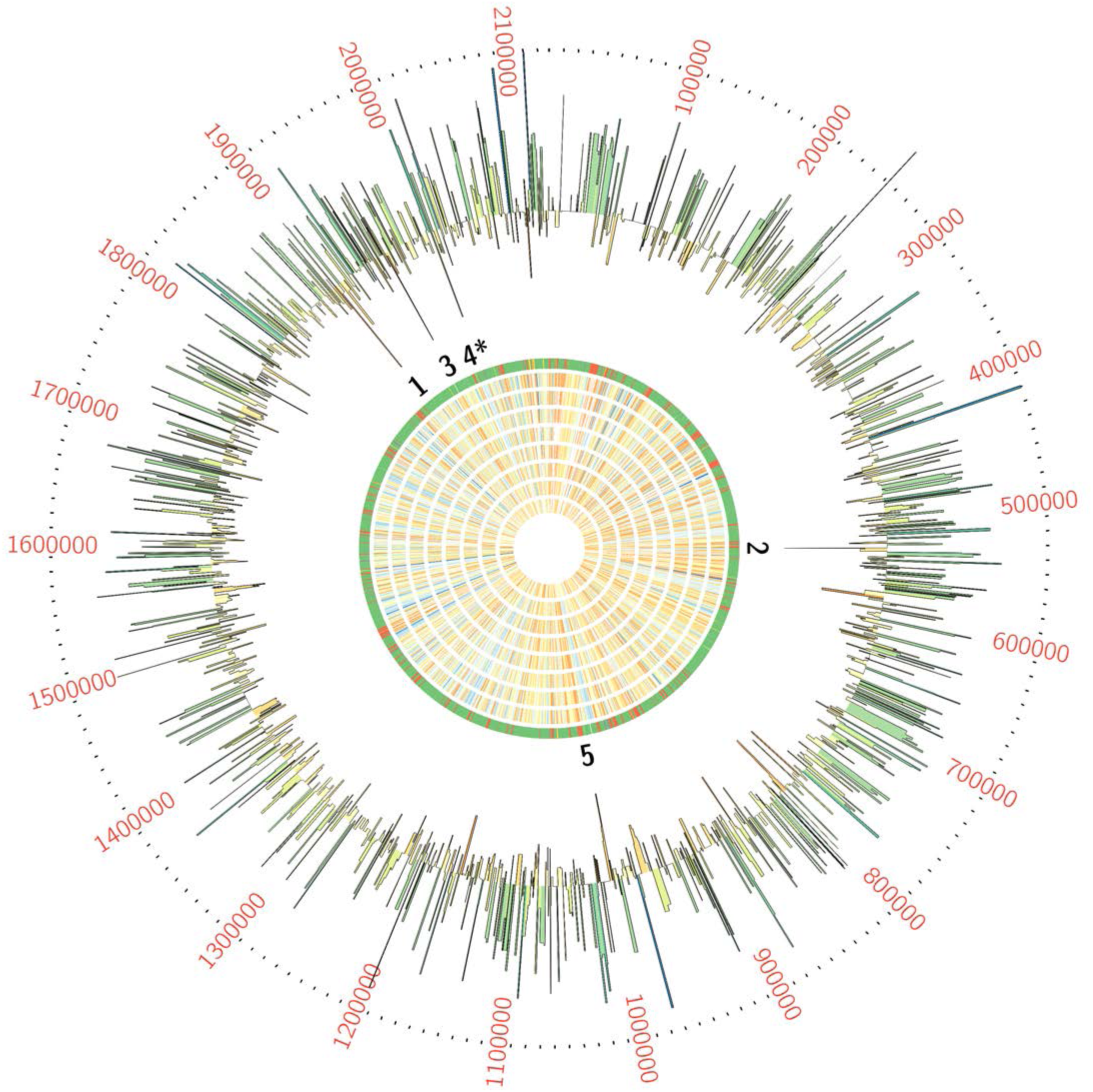
Tn-seq screening of GBS grown in human amniotic fluid revealed 5 conditionally essential genes. GBS transposon library in a background of strain A909 was grown in eight human amniotic fluid samples, then subjected to Tn-seq analysis. Each ring of the Circos plot shows a different representation of the A909 circular chromosome. From the center of the plot, rings 1-8 show transposon insertion density detected in the eight amniotic fluid outgrowth samples (lowest detection: red; highest detection: blue). Ring 9 shows essential genes (red), nonessential genes (green), intermediate genes (yellow), and those for which baseline fitness cannot be determined (white; see reference (41)). The outermost ring shows log_2_ fold-change values for each gene in amniotic fluid outgrowth as determined by ESSENTIALS. Genes with inward-pointing peaks had fewer than expected transposon insertions. Outward-pointing peaks had more than expected transposon insertions. The peaks are color coded for statistical significance (low adjusted p value: red; high adjusted p value: green). The five labeled peaks are described in the text and are listed in **Table 1**. Peak 4 is the GntR-class transcription factor, *mrvR*.

**Table 1:**
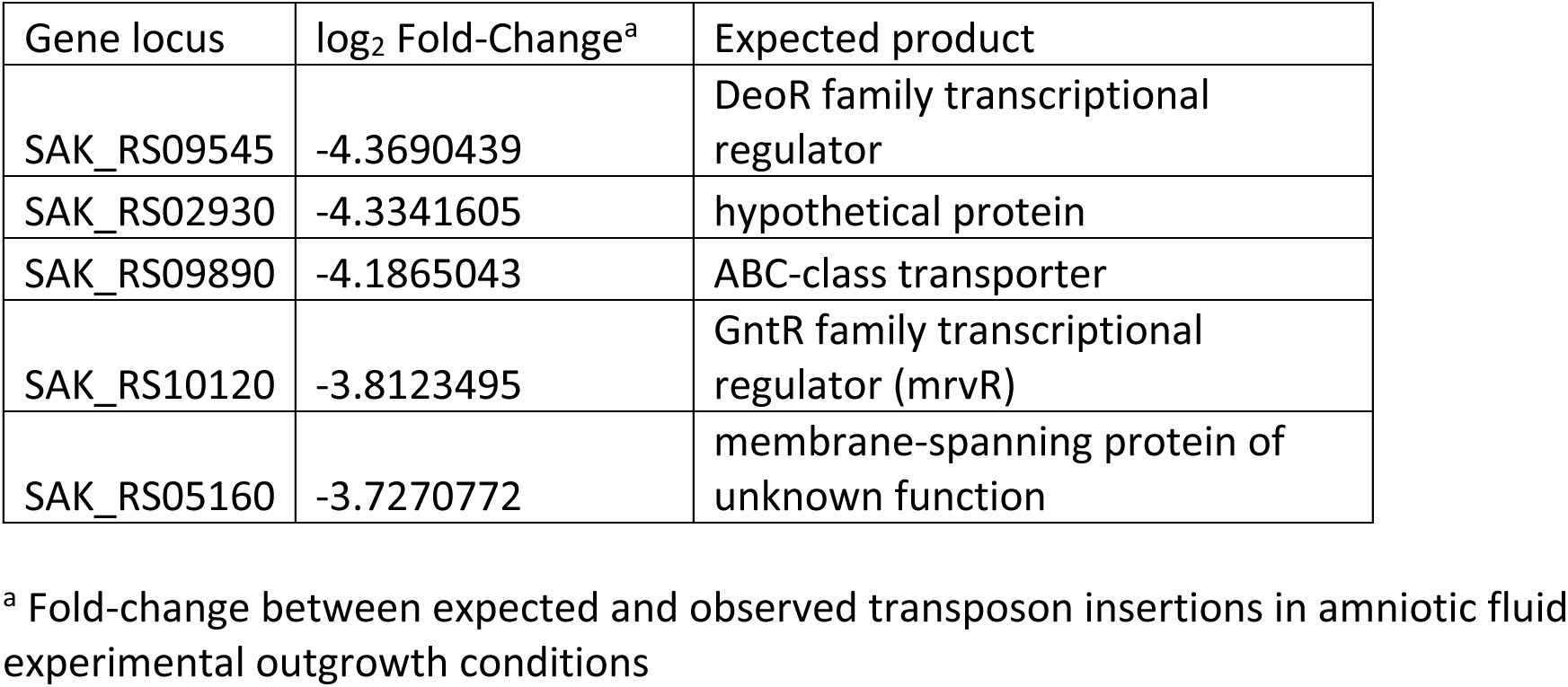
GBS genes conditionally essential for survival in human amniotic fluid

One of the candidate genes, SAK_RS10120, which we termed *mrvR*, had conserved functional domains suggesting its membership in the GntR superfamily of transcriptional regulators. The other candidate genes encoded an expected DeoR family transcriptional regulator (SAK_RS09545), a predicted membrane-spanning protein of unknown function (SAK_RS05160), an ABC-class transporter (SAK_RS09890), and one protein (SAK_RS02930) with a single domain of unknown function whose cellular localization and role are also unknown.

### CRISPRi knockdown and targeted knockout competition assays validate Tn-seq findings

To further validate the five candidate genes conditionally essential for GBS survival in amniotic fluid, we developed a CRISPRi gene expression knockdown system, which uses natively expressed, catalytically inactive Cas9 (dCas9) encoded on the GBS chromosome along with a single guide RNA (sgRNA) expression plasmid to generate efficient and flexible gene expression inhibition in GBS. We competed knockdowns of the five candidate conditionally essential genes against a sham targeting control dCas9 strain in human amniotic fluid and assayed survival using quantitative PCR analysis of knockdown and control strain persistence.

Development of a GBS CRISPRi system was aided by the fact that the GBS *cas9* gene shows high sequence similarity to the canonical *cas9* endonuclease first described in *Streptococcus pyogenes* (43–46) (**Supplemental Figure 1**). This homology allowed us to predict the two missense mutations that would establish a catalytically inactive isoform in GBS: D10A and H840A. The high level of homology between *S. pyogenes* Cas9 and the GBS homolog also suggested that the protospacer adjacent motif (PAM) necessary for Cas9 genomic target recognition likely followed the same NGG pattern as in *S. pyogenes*. We used a previously described sucrose-counterselectable mutagenesis plasmid, pMBsacB, to generate point mutations within the coding sequence of the *cas9* gene in GBS strain CNCTC 10/84 (serotype V) (47), which is the strain with which our animal models of vaginal colonization and ascending chorioamnionitis have been optimized.

We developed a sgRNA expression shuttle plasmid, p3015b, into which we cloned 20-bp protospacers against two genomic targets for each of the five candidate genes. We used the Broad Institute’s Genetic Perturbation Platform sgRNA design tool (https://portals.broadinstitute.org/gpp/public/analysis-tools/sgrna-design) to select optimized protospacers, then cloned synthetic oligonucleotide DNA into p3015b (48, 49). After confirming successful protospacer cloning, the targeting plasmids were purified and transformed into competent 10/84 modified to express dCas9.

To establish proof-of-concept for this approach we knocked down the first open reading frame (*cylX*) in the *cyl* operon that encodes genes responsible for synthesis of the pigmented cytotoxin *β*-hemolysin/cytolysin (50). p3015b encoding anti-*cylX* sgRNA could not be transformed into wild type GBS, presumably because Cas9 targeting to the chromosome resulted in lethal double-stranded DNA cleavage. The dCas9 strain, however, had high transformation efficiency and generated non-pigmented, non-hemolytic knockdowns when transformed with plasmid expressing anti-*cylX* sgRNA (**Figure 2A**).

**Figure 2.**
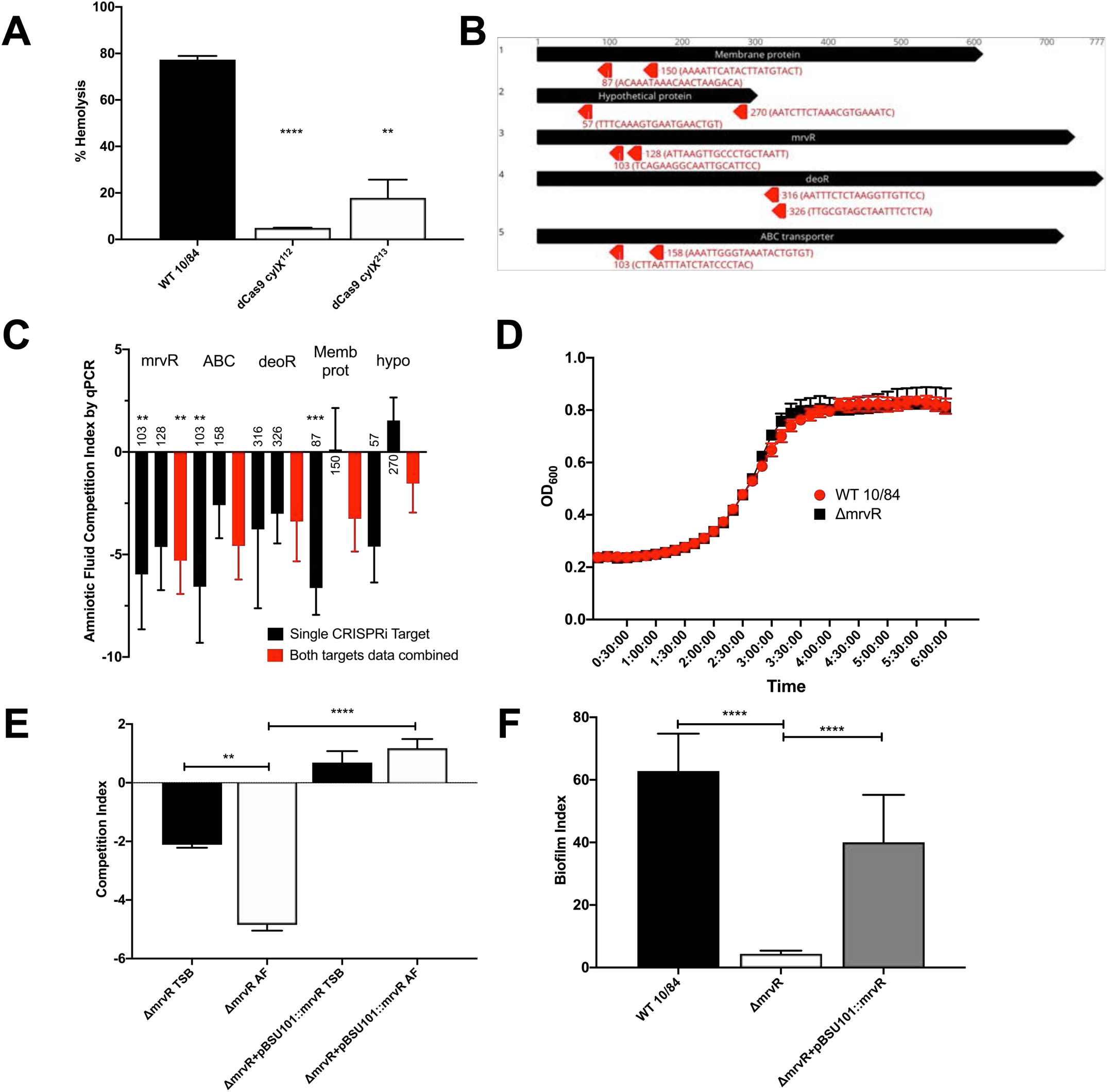
CRISPRi knockdowns of candidate genes and phenotypic characterization of GBS lacking MrvR in amniotic fluid, rich media, and biofilm-promoting growth conditions. Two CRISPRi targeting protospacers (targeting nucleotides 112 and 213 of *cylx*, the first gene in the operon) against the *cyl* operon were developed to knockdown expression of *β*- hemolysin/cytolysin. The resultant knockdown strains (in a 10/84 background) were compared to wild type in a human erythrocyte hemolysis assay (**A**). Targeting protospacers against the five candidate conditionally essential genes from amniotic fluid Tn-seq. For each gene, the number indicates the targeted nucleotide; the protospacer sequence is shown in parentheses (**B**). A qPCR-based multiplex CRISPRi screen using knockdowns of all five candidate genes revealed decreased fitness in amniotic fluid compared to tryptic soy (TS) broth (**C**); black columns show the competition index of strains bearing individual targeting protospacer plasmids, while red columns show pooled results from both targeting plasmids for each gene. Statistical significance of each knockdown phenotype was determined relative to growth of the knockdowns in TS broth. Growth curves of the *ΔmrvR* and wild type (WT) 10/84 in broth (**D**). Colony forming unit enumeration-based competition assays between *ΔmrvR* and *ΔmrvR*+pBSU101::*mrvR* in TS broth (TSB) and amniotic fluid (AF); **E**). Biofilm assay results comparing WT 10/84, *ΔmrvR*, and *ΔmrvR*+pBSU101::*mrvR*. *p<0.05, ** p<0.01, *** p<0.005, **** p<0.001; t test. Error bars show SEM surrounding mean.

Next, we created plasmids targeting two different sites in each of the five candidate amniotic fluid conditionally essential genes **(Figure 2B**) and transformed dCas9-expressing 10/84. As in previously described CRISPRi systems, we found optimal performance when the sgRNA targeting sequence matched sites within the open reading frame but on the non-template strand (51).

The resulting knockdown strains, plus a control dCas9 strain transformed with p3015b bearing a sham protospacer, were used to inoculate amniotic fluid and control broth. Plasmid DNA was purified at 24 hours of outgrowth. We used protospacer-specific quantitative PCR primers to characterize persistent GBS as targeting or non-targeting control. **Figure 2C** shows the results of these competition assays and demonstrates an expected competition defect in amniotic fluid for the five candidate knockdown strains.

As with CRISPRi knockdown systems in other organisms, the extent of the knockdown effect varied depending on the exact sequence targeted within the gene. In general, targets closer to the start codon showed more significant effects, which is consistent with other studies (51).

### A targeted knockout of mrvR shows a growth defect in human amniotic fluid

*MrvR* showed a consistent CRISPRi knockdown pattern, where both targeting protospacers generated a pronounced fitness defect in amniotic fluid. We selected this gene for further study based on these results and the hypothesis that this transcription factor may bridge microenvironmental changes associated with exposure to amniotic fluid and gene expression responses that promote amniotic fluid survival and possibly other GBS virulence traits.

Using pMBsacB, we generated an isogenic knockout of W903_RS09645, the *mrvR* ortholog in 10/84. We also complemented the gene deletion by cloning the full coding sequence of W903_RS09645 and its upstream promoter sequence into the shuttle vector pBSU101 (replacing the green fluorescent protein and *cfb* promoter sequence originally cloned into pBSU101) (52) and transforming the knockout with this plasmid.

We tested the knockout and complemented strains for survival defects in human amniotic fluid and in broth. **Figure 2D** shows growth curves of the wild type and knockout strains in tryptic soy broth, indicating that growth kinetics between the two strains in rich media are identical. **Figure 2E** shows results of competition assays in which amniotic fluid and broth were inoculated with a mixture of wild type 10/84 and the knockout strain or its complement. After 24 hours of outgrowth, these mixtures were plated for CFU enumeration on selective and nonselective media, permitting determination of the relative survival of the strains in competition.

We observed that in broth the knockout strain had a mild growth defect relative to wild type, but that the defect was more severe in amniotic fluid, and that both defects could be complemented through expression of the full-length *mrvR* gene.

### MrvR controls GBS biofilm formation

We observed phenotypic differences in growth of the *mrvR* knockout as compared to wild type 10/84. In liquid media, the mutant seemed to aggregate in clumps with less coating of the outgrowth vessel. We suspected that the knockout was biofilm deficient, which we confirmed using a biofilm assay. The biofilm index is the ratio of crystal-violet stained biofilm (following overnight growth in a multi-well plate) divided by the planktonic-phase optical density from the same sample. **Figure 2F** confirms that the knockout mutant is biofilm deficient compared to wild type 10/84, and that the biofilm phenotype is restored in the complemented strain.

### MrvR activity does not affect vaginal colonization

To test whether *mrvR* is necessary for persistent vaginal colonization, we colonized healthy adult female mice with knockout GBS or wild type control and performed serial surveillance swabs using an established protocol (13). We did not include the complemented knockout in this experiment because of past observations that, without ongoing *in vivo* antibiotic selection, plasmid DNA is rapidly cured by colonizing GBS.

While there was a trend toward earlier clearance among the mice colonized with the mutant strain, neither initial colonization density nor eventual clearance among the two groups differed significantly (**Figure 3**).

**Figure 3.**
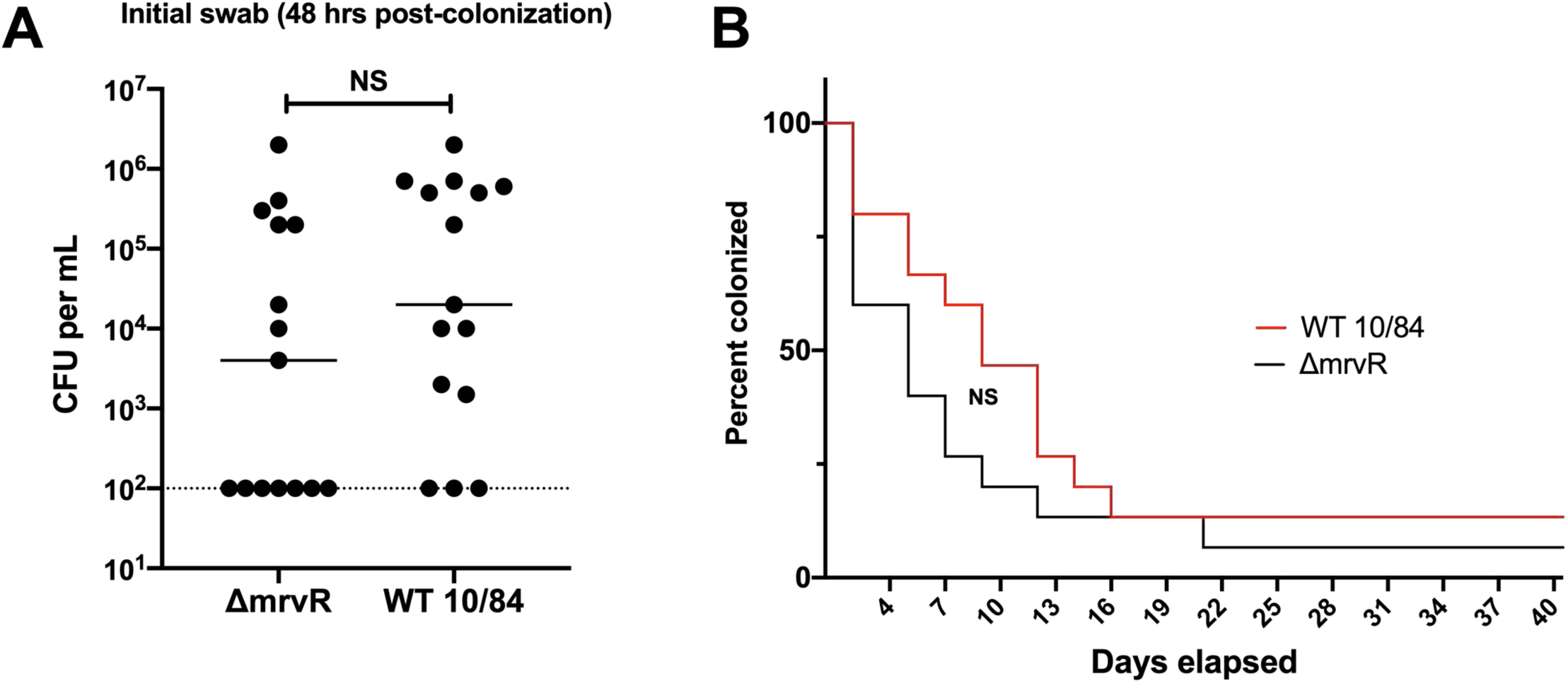
MrvR does not affect establishment or maintenance of vaginal colonization. Adult nonpregnant C57/BLJ mice were vaginally colonized with wild type (WT) 10/84 or *ΔmrvR*. Neither the colonization density at the initial swab 48 hours after inoculation (**A**) nor duration of colonization (**B**) showed significant differences (CFU: colony forming units, NS: not significant).

### The MrvR knockout causes chorioamnionitis without preterm delivery in a mouse model

To test the effect of *mrvR* in ascending chorioamnionitis, we used an established mouse model, vaginally colonizing pregnant female mice on pregnancy day 13 with either wild type 10/84 or the *mrvR* knockout.

Colonized pregnant mice were monitored through pregnancy day 17. Preterm delivery between day 13 and day 17 by a colonized mouse was one experimental endpoint, while animals that remained pregnant on day 17 were sacrificed and assessed for persistent vaginal colonization and ascending chorioamnionitis. Chorioamnionitis was established based on any of the following: GBS present in placental homogenates, GBS present in fetal homogenates, or GBS present in amniotic fluid. Data from mice that were no longer vaginally colonized on day 17 were not included in the analysis; none of the mice who cleared their vaginal colonization had chorioamnionitis.

There was a significant difference between pregnancy outcomes for mice colonized with wild type GBS and those colonized with the knockout. Whereas eight of eleven mice colonized with wild type GBS delivered preterm, none of the eleven mice colonized with the knockout strain did. However, all of the knockout colonized mice had chorioamnionitis with widespread GBS dissemination throughout the placentas, fetuses, and amniotic fluid. Of the mice colonized with wild type GBS, two of the three that carried their pregnancies through day 17 had chorioamnionitis (**Figure 4A-4B**).

**Figure 4.**
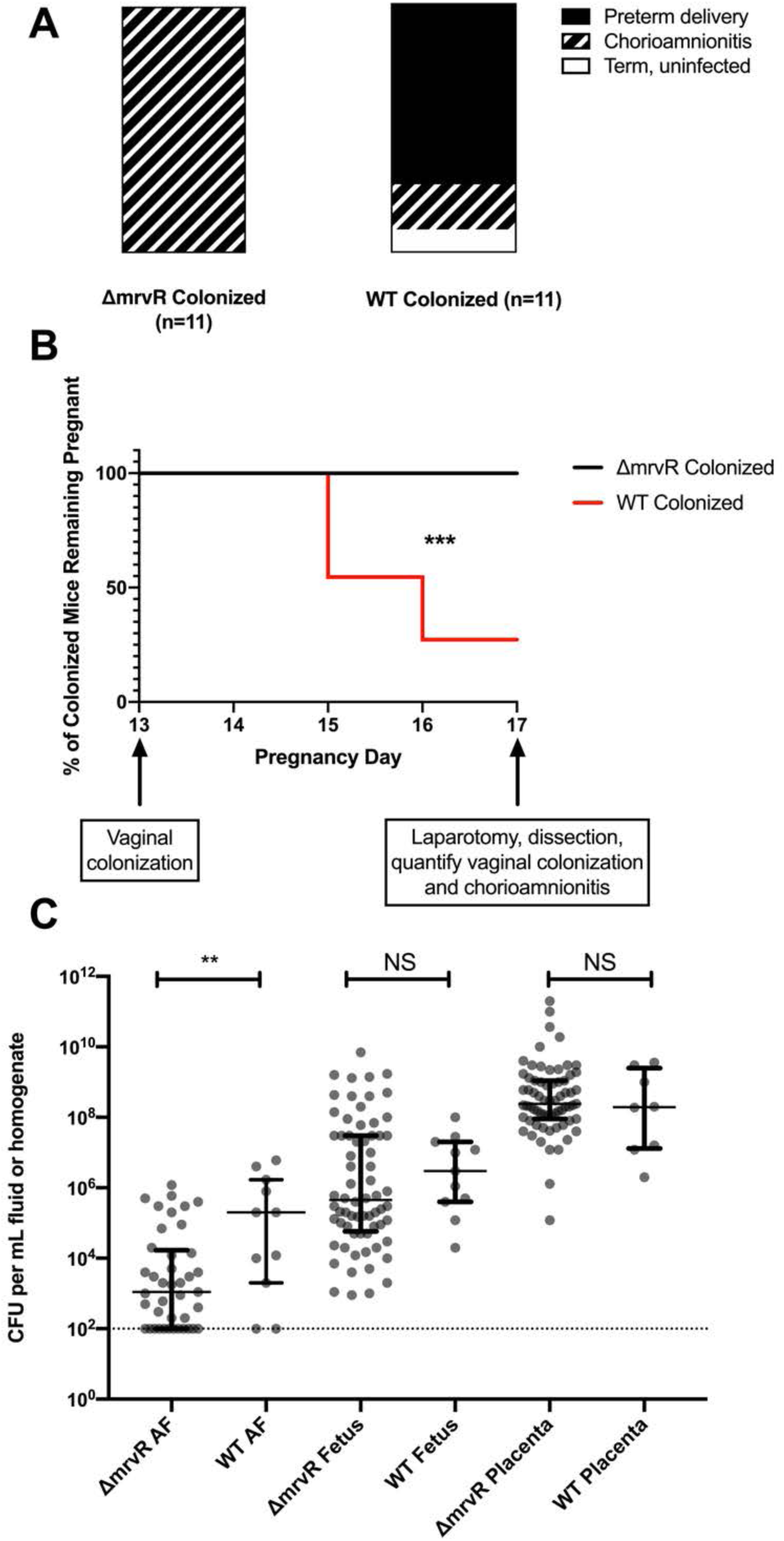
MrvR has a significant effect on the outcome of chorioamnionitis. Pregnant BALB/c mice were vaginally colonized with wild type (WT) or knockout (KO) 10/84 on pregnancy day 13. Pregnancies were monitored until day 17 or preterm delivery, whichever came first. On day 17, mice that remained pregnant were dissected and evaluated for vaginal colonization and chorioamnionitis. While none of the KO-colonized mice with chorioamnionitis delivered preterm, 8 of 11 WT-colonized mice delivered early. One WT-colonized mouse did not develop chorioamnionitis (**A-B**; *** p<0.005; Mantel-Cox test). Dissected pregnancy tissue was homogenized and quantified, showing a difference in invasion density in amniotic fluid (AF) but not in fetal or placental tissue (**C**, ** p<0.01; t test, error bars show interquartile ranges surrounding medians).

Colony counts from tissue and amniotic fluid samples collected at laparotomy showed a significant difference between wild type and knockout GBS density in amniotic fluid but not placental or fetal tissue. Bacterial density was higher in placental homogenates than in fetal tissue homogenates and amniotic fluid for both GBS strains, suggesting the placenta as the portal of infection (**Figure 4C**).

### MrvR is necessary for lethal invasive bacteremia in a murine sepsis model

To assess the role of the transcription factor in bloodstream invasion and systemic infection, we performed intraperitoneal injections of knockout and wild type GBS in healthy adult BALB-c mice. Both bacterial strains were transformed with a toxin-antitoxin stabilized plasmid expression system that generates constitutive luciferase, permitting *in vivo* monitoring of bacterial spread upon bacterial exposure to luciferin cofactor.

The infected mice were photographed using an IVIS *in vivo* imaging system at the time of initial intraperitoneal infection, then daily afterward until either death or complete clearance of the luciferase signal. Surviving mice that had cleared their infection based on imaging were subsequently observed for the following five days, and all remained well-appearing with normal behavior, eating, and stable weights throughout this post-imaging observation period.

This experiment revealed a significant difference in the ability of infected mice to clear mutant and wild type GBS. Whereas 87% of mice infected with wild type GBS died of the infection within three days, none of the mice infected with the knockout strain died. *In vivo* imaging showed that the mice were able to contain and clear the knockout strain, while the wild type strain tended to invade beyond the peritoneum such as into the thorax, a finding that was invariably followed by death (**Figure 5**).

**Figure 5.**
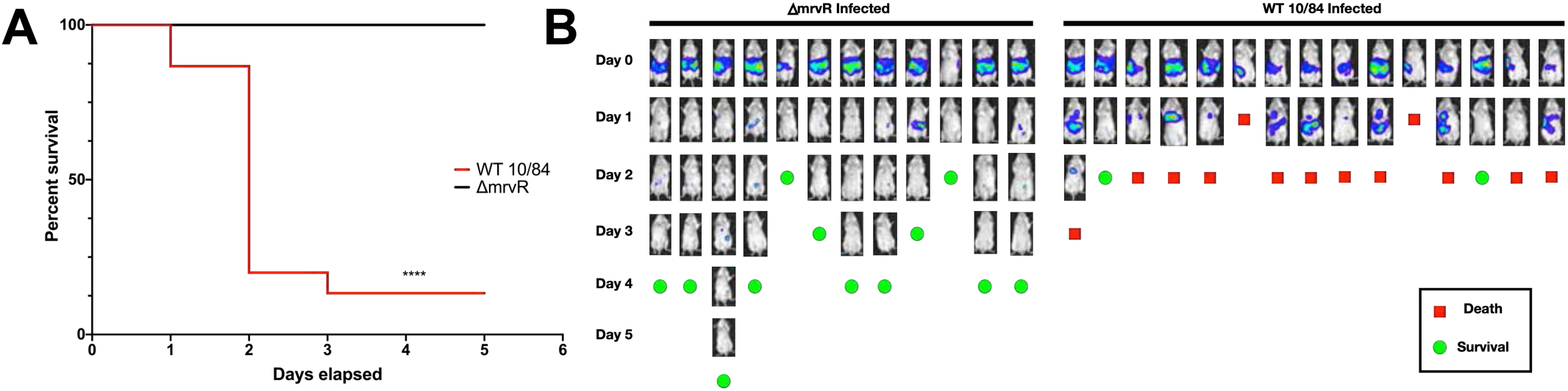
MrvR affects GBS invasiveness and lethality in a sepsis model. BALB/c mice were infected intraperitoneally with wild type (WT) 10/84 or *ΔmrvR*, both bearing a plasmid to allow in vivo tracking of a luciferase signal. There was a significant difference in outcome between the two strains (**A**, **** p<0.001 Mantel-Cox test) Mice were imaged daily (**B**) until they either died (red boxes) or completely cleared the luciferase signal (green circles). Mice that cleared the luciferase signal were monitored for the following five days and remained well.

**Figure 6.**
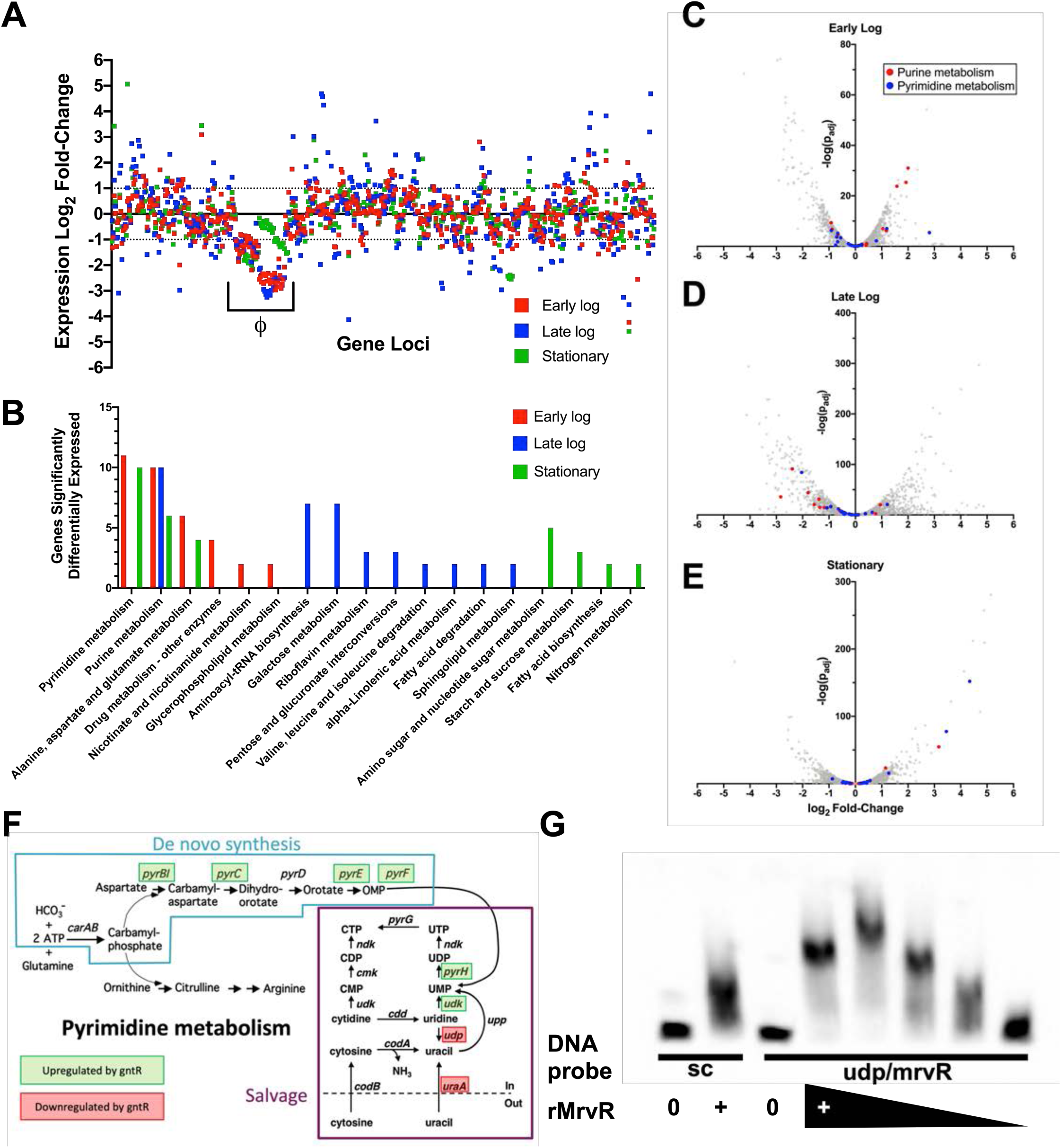
Analysis of the MrvR regulon reveals widespread effects with emphasis on genes involved in nucleotide metabolism. RNA-seq analysis of wild type (WT) and *ΔmrvR* 10/84 at three growth timepoints showed extensive regulation across the chromosome in all phases of growth (**A**, expression log_2_ fold-change shown relative to WT). A phage island (*Φ*) with significant differences from WT is noted. Gene set enrichment analysis showed that, across all three bacterial growth phases tested, the KEGG pathways most influenced by the presence of MrvR were pyrimidine and purine metabolism (**B**), which is also shown in volcano plots (**C-E**). Panel **F** shows specific genes of the de novo and salvage pathways of pyrimidine metabolism in GBS whose expression is significantly affected by the presence of MrvR. An electrophoresis mobility shift assay (EMSA) with recombinant MrvR (rMrvR) transcription factor combined with the shared promoter region between the *mrvR* and *udp* suggested increased protein binding (**G**) compared to a sequence scrambled control (sc). rMrvR had a maximal concentration of 20 mg/mL (+); in the experimental coincubation, each subsequent dilution was 2-fold in the same sterile PBS as the original suspension.

### Whole-genome transcriptomics shows that MrvR affects a widespread genetic network including multiple operons involved in nucleotide metabolism

We performed whole-genome RNA-seq on wild type GBS and the *mrvR* knockout in order to gain a preliminary understanding of the transcription factor’s regulon. We purified RNA from liquid cultures of the two strains at three growth time points: early log phase, late log phase, and stationary phase.

Comparing gene expression between the knockout and wild type GBS demonstrated that a substantial fraction of GBS transcription is controlled directly or indirectly by MrvR (**Figure 5A**). 16.8 percent of genes showed at least a 2-fold change in expression in the knockout relative to wild type during early logarithmic growth. This total increased to 35.7 percent in late logarithmic phase before dropping to 19.9 percent in stationary phase. Notably, a prophage island from W903_RS03075 through W903_RS03520 showed significant downregulation in the knockout at all three phases of growth. In order to confirm that this prophage DNA remained present in the knockout and had not undergone lysogeny, we performed PCR amplification of three prophage regions using genomic DNA template from the wild type and mutant GBS strains (**Supplemental Figure 2**).

We used a publicly available gene set enrichment search tool (53) to assess the differential gene expression data for functional patterns. This analysis revealed that genes involved in purine and pyrimidine metabolism were significantly enriched among the set of differentially expressed genes and that these nucleotide metabolism genes showed differential expression across all three growth phases. Other functional categories that showed significant differential expression tended to only be affected in one or, at most, two phases of growth (**Figure 5B-F**).

The *mrvR* gene is located next to a uridine phosphorylase gene (W903_RS09640), which encodes the Udp enzyme that converts uridine to uracil. The *udp* gene appears coregulated by MrvR, based on RNA-seq data. Intriguingly, the arrangement of MrvR adjacent to nucleotide metabolic genes— particularly uridine phosphorylase genes—is highly conserved across Gram-positive bacteria (**Supplemental Figure 3**).

### An electrophoresis mobility shift assay shows that the GntR transcription factor binds the intergenic region between its own coding sequence and the udp gene

Based on our transcriptomics data, we suspected that the transcription factor auto-regulates its own expression. To test this hypothesis, we used a *Brevibacillus*-based expression system to purify recombinant MrvR. We then used this recombinant sample to test DNA binding to the common promoter region between the coding region for MrvR and the adjacent *udp* gene using electrophoresis mobility shift assays. This experiment demonstrated a greater shift when the recombinant transcription factor was combined with the target DNA sequence as compared to a scrambled control DNA sequence with identical nucleotide ratios (**Figure 5G**), indicating autoregulation through binding of MrvR to its own promoter, which also coregulates *udp* expression.

## Discussion

GBS can survive and grow in diverse environments within the human host. It can persist in the intestine, in the male and female reproductive tracts, within the placenta, the amniotic fluid, human blood, joint spaces, the neonatal lung, and the central nervous system (1, 9, 12, 33, 54–58). This environmental tolerance contributes to GBS virulence, allowing it to invade and grow within compartments that are prohibitive to other bacteria.

This study began with a Tn-seq examination of genes that promote GBS fitness in amniotic fluid. GBS survival within human amniotic fluid contributes to its ability to cause ascending chorioamnionitis, which can lead to preterm labor and early-onset sepsis, sometimes in combination—a circumstance that can result in delivery of a severely infected preterm newborn with multi-organ system dysfunction (14, 59–62). Intraamniotic infection can also result in stillbirth (10). Amniotic fluid has chemical and immunological characteristics that are antimicrobial. The fetus and membranes produce antimicrobial peptides that inhibit bacterial survival, and amniotic fluid is intrinsically poor in nutrients required for bacterial proliferation (63–71).

We identified five GBS genes required for survival in human amniotic fluid. Two of these were predicted to encode transcriptional regulators, two encode surface-associated proteins, and one remains uncharacterized. A previous study that used a proteomic approach to evaluate differential gene expression in GBS strain A909 (the same background used for our Tn-seq library) during conditions associated with colonization and fetal invasion did not identify any of the same genes (72). Intriguingly, however, that investigation found two differentially expressed proteins (encoded by SAK_RS10135 and SAK_RS09535) whose genes are both two loci away from members of our set.

This study introduces a novel CRISPRi gene expression knockdown system, which we paired with qPCR to further interrogate the candidates suggested by our Tn-seq screen. Our approach builds on prior knockdown systems in other microorganisms and demonstrates some of the same findings, including the requirement that the DNA sequence targeted by the dCas9 enzyme must be on the antisense strand of the coding region and decreasing knockdown potency as the target region gets further from the start codon (51, 73, 74).

Our GBS CRISPRi system—which permits flexible and rapid phenotypic discovery without the challenge associated with generating gene deletions on the chromosome—could be used for diverse experimental purposes. In this case, it allowed us to validate Tn-seq results and to decide which candidate gene to further study. We envision additional uses, however, including rapid generation of knockdown libraries for multiplex testing of gene contributions to fitness in GBS and exploration of the roles of essential genes whose expression cannot be eliminated entirely.

In this study, the CRISPRi screen pointed our attention to a GntR-class transcription factor that seemed to play an important role in bacterial fitness in amniotic fluid. While GntR transcription factors are widely distributed throughout the bacterial kingdom, and some have been shown to play a role in virulence (75–83), this gene’s role in GBS host-pathogen interactions has not been explored. Interestingly, an orthologous gene was previously shown to influence group A *Streptococcus* susceptibility to the mouse cathelicidin CRAMP during murine skin infection (84). We investigated whether our knockout GBS had altered CRAMP sensitivity, but did not observe an effect. Instead we focused on various GBS phenotypic features that are regulated by this transcription factor.

We confirmed our CRISPRi knockdown findings with survival assays in broth and human amniotic fluid, using an in-frame deletion of the transcription factor, which we competed against the wild type parent strain. Unexpectedly, the MrvR knockout was visibly deficient at forming biofilms, which we subsequently confirmed with formal biofilm assays. GBS biofilms have been posited to influence colonization and host immune evasion in GBS and other bacteria (16, 85–87).

In a murine model of chorioamnionitis, absence of MrvR had a major impact on the ability of GBS to trigger preterm birth, with mice colonized on pregnancy day 13 showing consistent carriage of their pregnancies through day 17, despite extensive uterine invasion of the knockout strain. This is in contrast to the high rates of preterm delivery seen when pregnant mice are vaginally colonized with wild type GBS, which was observed in this study and in prior work (13) and has been documented in other animal models and clinical practice (14, 29).

This finding adds to accumulating evidence that GBS invasion of the fetoplacental unit is not, in itself, sufficient to trigger preterm delivery. A previous study demonstrated similar findings in an examination of the role of *β*-hemolysin/cytolysin in chorioamnionitis and preterm birth (13). Several others have shown that consequential inflammatory cascades are triggered in the choriodecidua following GBS invasion (14, 24, 30, 88). We hypothesize that these inflammatory cascades are enhanced by expression of virulence factors regulated by MrvR, such that its absence ameliorates the inflammation that leads to preterm birth without preventing tissue invasion; this hypothesis will be the focus of additional future work.

Our work also shows that pathways regulated by MrvR are critical for bloodstream invasion and host immune evasion. In an adult mouse model of sepsis, dramatic differences were seen between wild type and knockout GBS in terms of systemic spread and eventual mortality.

Our transcriptomic analysis of the *mrvR* knockout suggests that this transcription factor is involved in mediating nucleotide metabolism. Purine and pyrimidine sensing, biosynthesis, and salvage all play key roles in modulation of virulence traits in other bacterial pathogens (89–91). Furthermore, intracellular nucleotide homeostasis is highly regulated through diverse signaling pathways (89, 92, 93), lending plausibility to the prospect that the transcription factor under study here also participates in maintaining nucleotide concentrations. Our electrophoresis mobility shift assay results suggest that the transcription factor binds its own promoter region, which is also adjacent to the *udp* gene involved in purine metabolism—further suggesting a link between this signaling molecule and nucleotide availability. Future work will also focus on deeper characterization of the exact binding sites of this transcription factor across the GBS genome.

In conclusion, this study applied genome-wide screens, novel and established molecular techniques, and multiple murine models of GBS colonization and invasive disease to identify and characterize a GntR-class transcription factor, MrvR, with key roles in multiple virulence-related phenotypes. This newly described transcription factor presents an appealing subject for further mechanistic study and may be a target for novel pharmaceutical approaches to limit GBS virulence through modulation of this molecule’s activities.

## Methods

### Ethics statement

Animal experiments were performed under approved IACUC protocols at NYU and University of Pittsburgh. Adult phlebotomy for hemolysis assays was conducted under a University of Pittsburgh approved IRB protocol. Collection of anonymous, discarded amniotic fluid was conducted under an approved exemption from the NYU IRB.

### Statistical analyses

Statistics for Tn-seq were determined with ESSENTIALS software (94) and RNA-seq statistical tests were performed with DESeq (95). Remaining statistics for figures were calculated using Prism for macOS v. 8.4.3 (Graphpad Software, San Diego, CA).

### Bacterial strains and growth conditions

GBS strains A909 (serotype Ia, sequence type 7) and CNCTC 10/84 (serotype V, sequence type 26) and their derivatives were grown at 37°C (or 28°C when the temperature-sensitive pMBsacB plasmid was present and extrachromosomal) under stationary conditions in tryptic soy (TS) medium (Fisher Scientific cat. # DF0370-17-3), supplemented with 5 µg/ml erythromycin, 150 µg/ml spectinomycin, or rifampin 10 µg/ml as needed for selection. *Escherichia coli* was grown at 37°C (or 28°C with extrachromosomal pMBsacB present) with shaking in Luria-Bertani (LB) medium (Fisher Scientific cat. # DF9446-07-5) supplemented with 300 µg/ml erythromycin and 150 µg/ml spectinomycin as needed for selection. *Brevibacillus choshinensis* for expression of recombinant MrvR was grown at 28°C in TM or 2SY media supplemented with neomycin 50 μg/mL according to the recommendations of Takara Bio.

### Tn-seq on GBS grown in human amniotic fluid

Discarded, anonymous amniotic fluid from amniocentesis was obtained from the NYU-Langone Medical Center Department of Obstetrics and Gynecology. Samples were stored frozen at - 20°C, then thawed on ice. Before use in outgrowth experiments, the amniotic fluid was passed through a 0.2 µm sterile filter.

A previously described saturated transposon mutant library in an A909 background (41) was thawed from frozen stock. The library was washed twice in sterile PBS to remove glycerol cryoprotectant, then resuspended in 3 mL PBS and seeded into 100 mL of TS broth with erythromycin selection. After overnight outgrowth, 20 mL of the library was seeded into fresh TS broth with erythromycin and allowed to grow to mid-log phase (OD_600_=1). 120 mL of this culture was then pelleted and washed twice with PBS, then resuspended in 1 mL of PBS.

50 µL of this concentrated library was then used to seed either 5 mL of amniotic fluid pre-warmed to 37°C or control TS broth. The seeded amniotic fluid and TS was allowed to grow overnight. Unseeded control amniotic fluid samples were also incubated and plated the following morning to ensure that there was no contamination present.

After overnight outgrowth, the TS and amniotic fluid samples were transferred into 250 mL aliquots of TS. These were grown overnight, then genomic DNA was purified from 90 mL of each culture using the MoBio Powersoil DNA Extraction Kit (cat. # 128888) according to manufacturer instructions. Purified DNA was subsequently digested with MmeI and used for barcoded terminal adapter ligation, PCR, and sequencing on a 150-nt paired-end run of the Illumina (San Diego, CA, USA) HiSeq 4000 platform, with a target number of reads per library of 50 million and subsequent demultiplexing as previously described (41, 42).

We used ESSENTIALS, a publicly available Tn-seq analysis server, to analyze our data (94). ESSENTIALS generates a plot of genome-wide log_2_ fold-change values for actual-versus-expected transposon detection. Bimodality in this plot indicates separate gene populations— those with approximately the expected number of transposon insertions detected in the experimental condition, and those with fewer than expected detected insertions. The latter group represents the set of conditionally essential genes, and the local minimum between the two modal peaks can be used to generate a cutoff value between nonessential and conditionally essential genes. In the case of our amniotic fluid Tn-seq experiment, the local minimum was at a log_2_ fold-change value of -3.22. The five candidate genes evaluated had log_2_ fold-change values below this cutoff (**Supplemental Data 1**).

### Development of a GBS CRISPRi system for gene expression knockdown

To generate GBS expressing dCas9, we used the mutagenesis plasmid pMBsacB in GBS strain 10/84 as previously described (47). Two separate synthetic, double-stranded DNA fragments, 800 to 1000 nt in length, were designed and ordered from Genscript (Piscataway, NJ, USA). The fragments matched the native 10/84 *cas9* coding sequence except for the two missense mutations, D10A and H840A, needed to render the enzyme catalytically inactive. These fragments were cloned into pMBsacB using Gibson assembly and used to transform chemically competent DH5*α E. coli* (New England Biolabs, Ipswich, MA, USA) according to manufacturer instructions. We confirmed proper cloning by Sanger sequencing of miniprepped plasmid DNA, which we then used to transform electrocompetent GBS as previously described (47, 96, 97). We performed the C-terminal (H840A) mutagenesis first, following the protocol we have described previously (47). Once this mutation was generated and confirmed by Sanger sequencing, we made those GBS competent and created the second mutation using identical techniques.

p3015b is a shuttle vector originally derived from pVPL3004 (98). We used PCR, synthetic DNA, and Gibson assembly approaches to make multiple changes to this plasmid for use in GBS. The kanamycin resistance cassette was changed to the erythromycin resistance gene; the *cas9* gene from *S. pyogenes* was removed; and the tracrRNA sequence downstream of the dual BsaI restriction sites used for protospacer introduction was replaced with a new sequence that conforms to the predicted folding structure of the native GBS tracrRNA sequence (**Supplemental Figure 1**). We also sought to introduce anhydrotetracycline inducibility of sgRNA expression by cloning a *tetR* coding sequence and a 2x tetO inducible promoter upstream of the sgRNA coding sequence. Despite extensive efforts, however, we could not suppress baseline sgRNA expression in GBS to the point that gene knockdown was inducible (data not shown), so the plasmid was used for constitutive sgRNA expression in GBS without anhydrotetracycline addition.

Targeting protospacers were designed using the Broad Institute’s online sgRNA design tool (48, 49). Target sequences were selected based on their antisense orientation and, where possible, position in the first half of the coding sequence. Protospacer oligonucleotide sequences were ordered from Integrated DNA Technologies (Coralville, IA, USA) and cloned into BsaI-digested 3015b as originally described by Jiang et al. in their description of pCas9, from which pVPL3004 was developed (99).

### CRISPRi multiplex competition assay

Cultures of 10/84 dCas9 transformed with p3015b plasmid bearing targeting protospacers against candidate conditionally essential genes were grown overnight in selective broth. A control strain with p3015b plasmid containing a sham protospacer with no matching sequence in the GBS chromosome was grown as a control. The 11 cultures were spun down and washed twice in PBS, then resuspended and normalized to an OD_600_ of 0.8. These suspensions were then plated as serial dilutions to confirm that the starting CFU concentrations were approximately equal.

500 μL from each of the 11 normalized GBS suspensions were then combined to make a starting mixed culture. A sample of this mixed culture was used for plasmid DNA purification using the Thermo MagJET plasmid DNA Kit (cat. # K2791) according to manufacturer instructions.

Amniotic fluid or TS broth containing erythromycin for continued plasmid selection was pre-warmed to 37°C and seeded 1:50 with the mixed culture, then allowed to grow stationary overnight. 24 hours after seeding, plasmid DNA was extracted from the amniotic fluid and TS samples.

At both the starting and 24-hour timepoints, prior to purification, DNA from dead bacteria was removed using ethidium monoazide using techniques described by Soejima et al. (100).

Ethidium monoazide—which binds to extracellular DNA or DNA within dead bacteria, but not to DNA within live cells—was added to a final concentration of 10 μg/mL, after which the cultures were allowed to incubate at 4°C for five minutes. They were then placed under a high intensity white light source, which destroys DNA bound to ethidium monoazide. The treated samples were then spun down, washed once in PBS, then used for plasmid purification.

qPCR on purified plasmid samples from the initial and 24-hour timepoints was performed using Applied Biosystems Power SYBR Green Master Mix (Thermo cat. #4368577) on a BioRad (Hercules, CA, USA) CFX384 real-time PCR thermocycler. The different plasmids present were detected using a conserved reverse primer and a forward primer complementary to either one of the targeting protospacer sequences or the sham protospacer sequence. We also used a pair of normalization primers complementary to plasmid sequences distant from the protospacer cloning site. This strategy allowed determination of relative abundances of each of the targeting protospacers (and the sham control) relative to the total plasmid concentration in each sample.

Competition indices for each knockdown strain were calculated using the formula: ln((NE^target^_T24_/NE^target^_T0_)/(NE^sham^_T24_/NE^sham^_T0_)) where NE is normalized expression scaled to total plasmid quantity, target and sham reflect targeting and control plasmids, and T0 and T24 represent samples from the initial mixed culture or from the 24-hour timepoint, respectively. qPCR was performed twice, with two to three technical replicates per primer pair. Competition indices for the knockdown strains were normalized to survival for each strain in TS broth.

### Generation of the mrvR knockout and its complemented control strain

A synthetic double-stranded DNA fragment was designed to contain the chloramphenicol acetyltransferase gene surrounded by 500-nt homology arms that match upstream and downstream sequences flanking the *mrvR* open reading frame in 10/84. This mutagenesis cassette was cloned into pMBsacB using Gibson assembly, then used to transform electrocompetent wild type 10/84. Temperature- and sucrose-based selection and counterselection were used to isolate single-cross and double-cross plasmid insertion and excision mutants, which were confirmed using targeted PCR and Sanger sequencing as previously described (47).

To complement the gene deletion, the wild type 10/84 *mrvR* gene and 75 nt of upstream promoter sequence were amplified using PCR and cloned into pBSU101 in place of the *cfb* promoter and green fluorescent protein gene present in the original plasmid (52). This plasmid was then used to transform the knockout strain, using spectinomycin for positive selection.

### Amniotic fluid competition assays with CFU quantification

Amniotic fluid or TS broth was seeded with either a) a 10/84 strain with a defined mutation in the *rpoB* gene that confers rifampin resistance (101) and a spontaneous streptomycin-resistant knockout clone or b) the rifampin-resistant 10/84 strain transformed with empty pBSU101 (for spectinomycin resistance) and the complemented, streptomycin-resistant knockout strain. After 24 hours of outgrowth, these mixtures were plated for CFU enumeration on nonselective media, rifampin-containing media, and streptomycin-containing media. This permitted determination of total growth and allowed discrimination between the knockout and wild type strains.

Competition indices were calculated using colony forming unit (CFU) density per mL, where competition index=ln((CFU^expt^_T24_/CFU^expt^_T0_)/ (CFU^WT^_T24_/CFU^WT^_T0_)); here, expt represents the experimental strain (either knockout or control) and WT is wild type 10/84. Assays were performed in triplicate and the experiment was repeated twice.

### Hemolysis assays

Whole cell GBS was used in hemolysis assays on washed human erythrocytes as previously described (42).

### Growth curve analysis

Wild type and *mrvR* knockout GBS were grown to late logarithmic phase in TS broth. The two cultures were then normalized to an OD_600_ of 1.0, then diluted 1:50 in broth. Triplicate samples were then instilled into a clear, flat-bottom 96-well plate, the lid of which had been treated with sterile defogging solution. OD_600_ absorbance readings were taken every 10 minutes, with a brief shaking step before each read, using a Molecular Devices (San Jose, CA, USA) SpectraMax M4 plate reader set to 37°C.

### Biofilm assays

10 μL of stationary-phase GBS was used to seed 1 mL of fresh broth with appropriate antibiotic selection and 5% glucose supplementation in a 12-well, clear-bottom plate, which was then allowed to grow for 24 hours at 37°C. The plate was rocked gently on an orbital shaker for 1 minute to resuspend non-adherent bacteria, then the media was carefully removed and transferred to a fresh plate. OD_600_ absorbance readings of the planktonic phase bacteria were recorded.

The plate that grew overnight was then used to stain the residual biofilms. 500 μL of 1% crystal violet was carefully added to each well, being cautious not to disturb the biofilm. After 10 minutes of gentle shaking at room temperature, the stain was removed and each well was washed three times in 1 mL of sterile PBS. After the final wash, the stained biofilm was allowed to dry, then solubilized with 1 mL of a 20% acetone/80% ethanol solution. Absorbance at 550 nm was measured. Biofilm indices were calculated as OD_550_/OD_600_. The experiment was performed three times with three technical replicates for each condition.

### Murine vaginal colonization model

Vaginal colonization of nonpregnant C57BL/6J mice was performed as previously described by Randis et al. (13) with minor modifications.

6- to 8-week old mice were injected subcutaneously with 0.5 mg *β*-estradiol on two successive days to synchronize estrus. On the third day, stationary phase GBS was pelleted and resuspended in a 1:1 mixture of PBS and a sterile 10% gelatin solution to enhance viscosity. The mixtures were diluted and plated for CFU enumeration. Mice were vaginally colonized with 50 µL, corresponding to 10^7^ CFU.

Following colonization, mice were single-housed. After 48 hours, an initial swab to detect colonization was performed. A sterile nasopharyngeal swab was moistened in 300 µL sterile PBS, then inserted into the murine vagina and rotated three times, after which it was replaced in the PBS and swirled to release adherent GBS. The PBS was diluted and plated for CFU enumeration on GBS-specific Chromagar plates (Chromagar, Paris, France, cat. # SB282), which were then grown overnight at 37°C.

Mice with established vaginal colonization, based on the 48-hour swab, were then swabbed three times weekly (Monday, Wednesday, and Friday) following the above protocol. Colonization clearance was defined as two sequential swabs with no detectable GBS.

### Chorioamnionitis and preterm delivery model

The murine chorioamnionitis/early-onset sepsis model was performed as previously described (13) with minor modifications. Two- to four-month old BALB/c mice were mated and plug-positive female mice were monitored for pregnancy by physical examination and serial weights.

On pregnancy day 13, mice were vaginally colonized with either wild type 10/84 or the *mrvR* knockout as in the vaginal colonization model (without the preceding estradiol injections). Following colonization, they were single-housed and monitored daily for preterm delivery. If delivery occurred prior to day 17, this outcome was considered chorioamnionitis with preterm birth. Mice that remained pregnant on day 17 were sacrificed and dissected. Each fetus was examined for signs of intrauterine fetal demise (significant pallor or hyperemia, visible skin sloughing or purulence, or evidence of ongoing fetal resorption). An insulin needle was used to withdraw amniotic fluid from within the fetal membranes, after which placentas and fetuses were isolated and homogenized.

Amniotic fluid and tissue homogenates were diluted and plated on Chromagar plates for CFU enumeration, which was performed after overnight plate growth at 37°C.

### Generation of luciferase reporter GBS

A toxin-antitoxin stabilized shuttle vector expressing the firefly luciferase gene, pLZ12Km2-P23R:TA:ffluc, was a gift from Thomas Proft (Addgene plasmid # 88900 ; http://n2t.net/addgene:88900 ; RRID:Addgene_88900). We used Gibson assembly techniques to excise the kanamycin resistance marker on pLZ12Km2-P23R:TA:Ffluc and replace it with an erythromycin resistance gene. The new plasmid, called pFfluc-Erm, was purified from *E. coli* and used to transform electrocompetent wild type 10/84 and the *mrvR* knockout.

In order to fluoresce, GBS bearing the pFfluc-Erm plasmid must be exposed to the cofactor D-luciferin (Millipore Sigma cat. # L9504), which was prepared in a filter-sterilized stock solution at a concentration of 28 mg/mL in water. We performed *in vitro* imaging of serial dilutions of liquid cultures of GBS transformed with pFfluc-Erm combined 1:1 with D-luciferin stock and determined the level of fluorescence detection at 290,000 CFU/mL.

### Adult mouse sepsis model with in vivo imaging

Six- to eight-week old female BALB/c mice were injected intraperitoneally under general anesthesia with 10^8^ CFU of PBS-washed, stationary phase WT 10/84 or *mrvR* knockout 10/84 bearing the pFfluc-Erm plasmid and 500 µg of D-luciferin in 500 µL PBS. Immediately after the injection, mice were imaged on a Perkin-Elmer (Waltham, MA, USA) IVIS Lumina Series II imaging instrument with a 1-minute exposure time.

On subsequent days, surviving mice were injected intraperitoneally with 560 µg D-luciferin in sterile water then imaged after 10-15 minutes using a 1-minute exposure time. Mice that cleared their luciferase signal were subsequently observed for five days.

### Whole-genome RNA-seq transcriptomic analyses

Wild type 10/84 and the *mrvR* knockout 10/84 strain were grown in TS broth overnight. 13-mL overnight culture samples were used for stationary phase RNA extraction. This culture was also used to seed pre-warmed broth, which was then grown to an OD_600_ 0.3-0.6 (early log phase) or 0.9-1.1 (late log phase), at which point RNA extraction was performed.

We used the Ribopure Bacteria RNA Purification Kit (Thermo cat. # AM1925) according to manufacturer specifications with the following exceptions: the initial bead beating step was performed for 20 minutes; the early log bacterial RNA was extracted from an 85-mL starting culture while late log and stationary phases were extracted from 13-mL cultures. RNA samples were then treated with the DNase and DNase inactivation reagents supplied with the kit.

Sequencing library preparation was conducted with the Illumina TruSeq total RNA kit according to manufacturer instructions. rRNA depletion was performed using the Ribo-Zero Plus rRNA Depletion Illumina. Next, random primers initiated first and second strand cDNA synthesis.

Adenylation of 3’ ends was followed by adapter ligation and library amplification with indexing. Sequencing was performed on an Illumina NextSeq500 platform with paired-end 75-nt reads over 150 cycles.

Trimmed and demultiplexed reads were aligned to the 10/84 genome using Bowtie 2 (102). These alignments were then used for statistical analysis using DESeq (95), which allowed determination of expression fold-change and p values. Gene set enrichment analyses were performed with Genome2D (53).

### Purification of recombinant MrvR

The coding sequence for MrvR was PCR amplified from 10/84 genomic DNA using primers optimized for subsequent cloning into the four supplied BIC *Brevibacillus* expression system pBIC plasmids (Takara cat. # HB300/HB310) in order to generate secreted, N-terminus poly-His tagged recombinant MrvR. PCR, cloning, and transformation steps were carried out in accordance with manufacturer recommendations. Resultant colonies were PCR screened and Sanger sequenced for successful cloning. Optimal secretion of recombinant protein was observed in *Brevibacillus* bearing the pBIC3 plasmid construct. This strain was then grown in 2SY medium supplemented with neomycin for 72 hours at 28 °C.

His-tag affinity purification was performed with Takara Capturem maxiprep columns (cat. # 635713) according to manufacturer instructions. Following protein elution through the column, the product was concentrated and buffer exchanged into 1x PBS pH 7.4 using a Pierce protein concentrator column (Thermo cat. # 88527).

SDS-PAGE was performed under denaturing conditions using Thermo Scientific Bolt 4-12% Bis-Tris precast gels (cat. # NW04120BOX) and recommended reagents. For Coomassie staining, purified His-tagged MrvR was analyzed along with unprocessed supernatant before and after passage through a 0.2 μm filter to remove any planktonic cells, flow-through, and the final membrane wash flowthrough (**Supplemental Figure 4A**). For Western blot, three separate purifications of MrvR were run with a His-tagged human CD4 protein as a positive control (**Supplemental Figure 4B**). Following transfer to a PVDF membrane, Rabbit anti-His tag monoclonal IgG (Fisher cat. # TA591008S) was used to probe the blot, followed by chemiluminescent detection using the Pierce Fast Western Blotting Kit with SuperSignal Rabbit West Pico reagents (Thermo cat. #35061).

### Electrophoresis mobility shift assay

Electrophoresis mobility shift assay was performed with the Thermo Lightshift Chemiluminescent EMSA Kit (cat. # 20148) according to manufacturer instructions with the following details. The 153-nt *mrvR* and *udp* promoter region was PCR amplified using a biotinylated forward primer and gel purified. A scrambled biotinylated DNA fragment of the same length and with the same nucleotide ratios was used as a control. Mixtures of undiluted, purified recombinant MrvR (20 mg/mL) and four 2-fold dilutions were combined with promoter DNA at a fixed concentration, while only undiluted rMrvR was combined with the scrambled control fragment. All reactions included poly (dI•dC) as a nonspecific blocking reagent, as well as 1x binding buffer included with the kit. Additional negative controls were performed in which no rMrvR was present. Coincubation continued for 20 minutes at room temperature, after which the reactions were run on a 4-6% polyacrylamide gel under native conditions. Blotting and detection of the labeled DNA fragments was performed in accordance with instructions with the EMSA kit.

## Acknowledgements

We are grateful to the University of Pittsburgh Center for Research Computing, which maintains the HTC cluster used for bioinformatic analysis of RNA-seq data. William MacDonald and Rania Elbakri in the University of Pittsburgh Health Sciences Sequencing Center assisted with next-generation sequencing of RNA-seq samples. Staff at the University of Maryland Genomic Resource Center assisted with sequencing for Tn-seq. Thomas Proft kindly made the plasmid pLZ12Km2-P23R:TA:ffluc available through Addgene. We thank Thomas Diacovo and Jeffrey Weiser, who shared equipment for multiple experiments in this study.

## Funding

This work was supported by NIH/NIAID grants K08AI132555 to T.A.H., R21AI147511 to T.A.H. and A.J.R., and R01AI143290 to A.J.R.; the Pittsburgh Children’s Foundation Children’s Trust Young Investigator Award (T.A.H.); the Richard King Mellon Institute for Pediatric Research Pilot Award (T.A.H.); and the i4Kids Pilot Award (T.A.H.).

## Data availability

Tn-seq and RNA-seq reads available on SRA.

## Supplemental Data Files

**Supplemental Data 1: GBS amniotic fluid conditionally essential gene analysis** Output from ESSENTIALS analysis of eight A909 library aliquots grown in human amniotic fluid. The table is sorted by log_2_ fold-change (D), and the five conditionally essential genes described in the main text are highlighted in yellow. The index (column A) allows sorting by gene locus if set from smallest to largest.

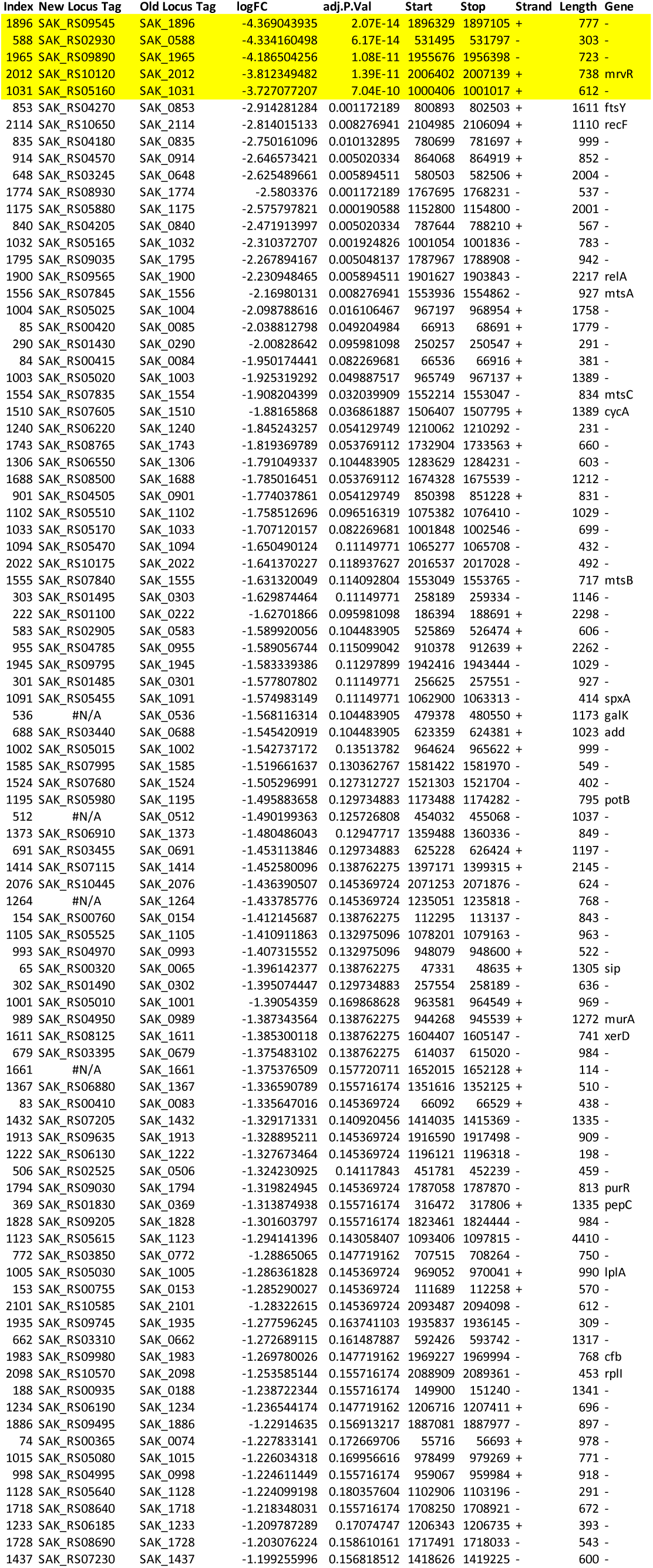
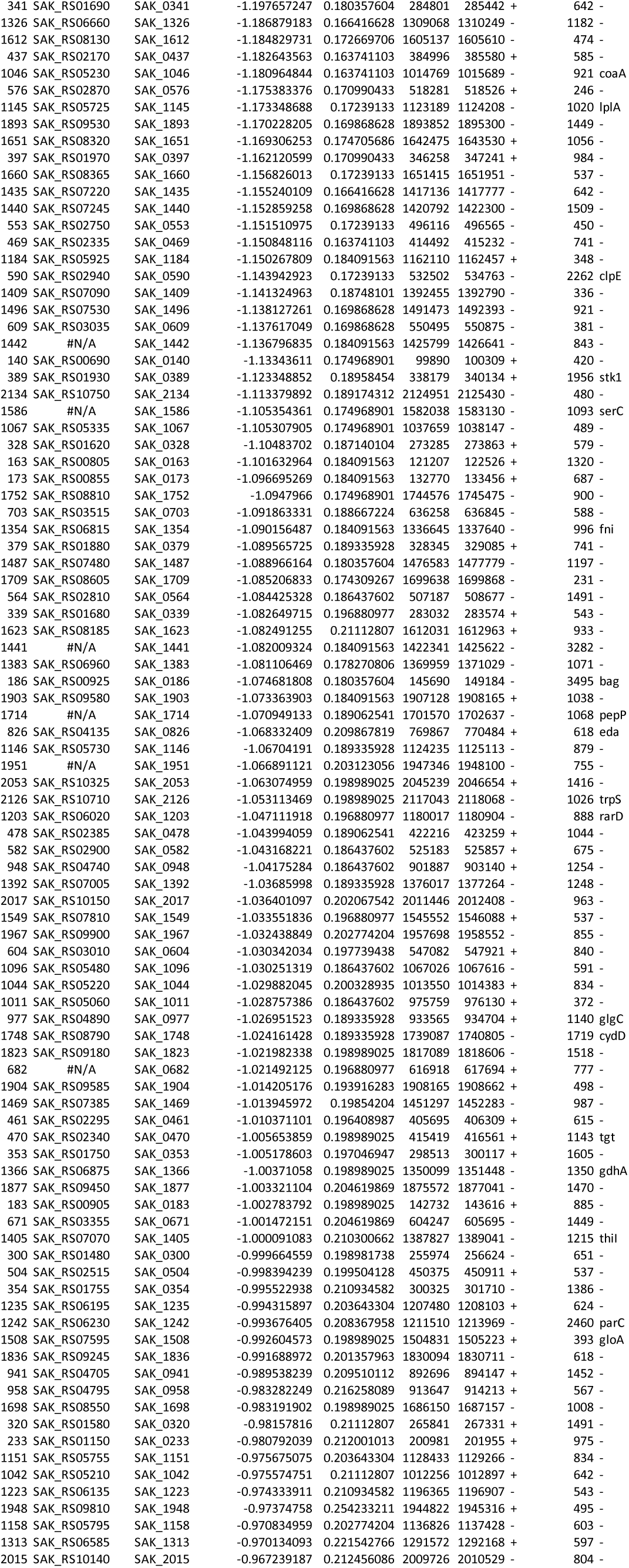
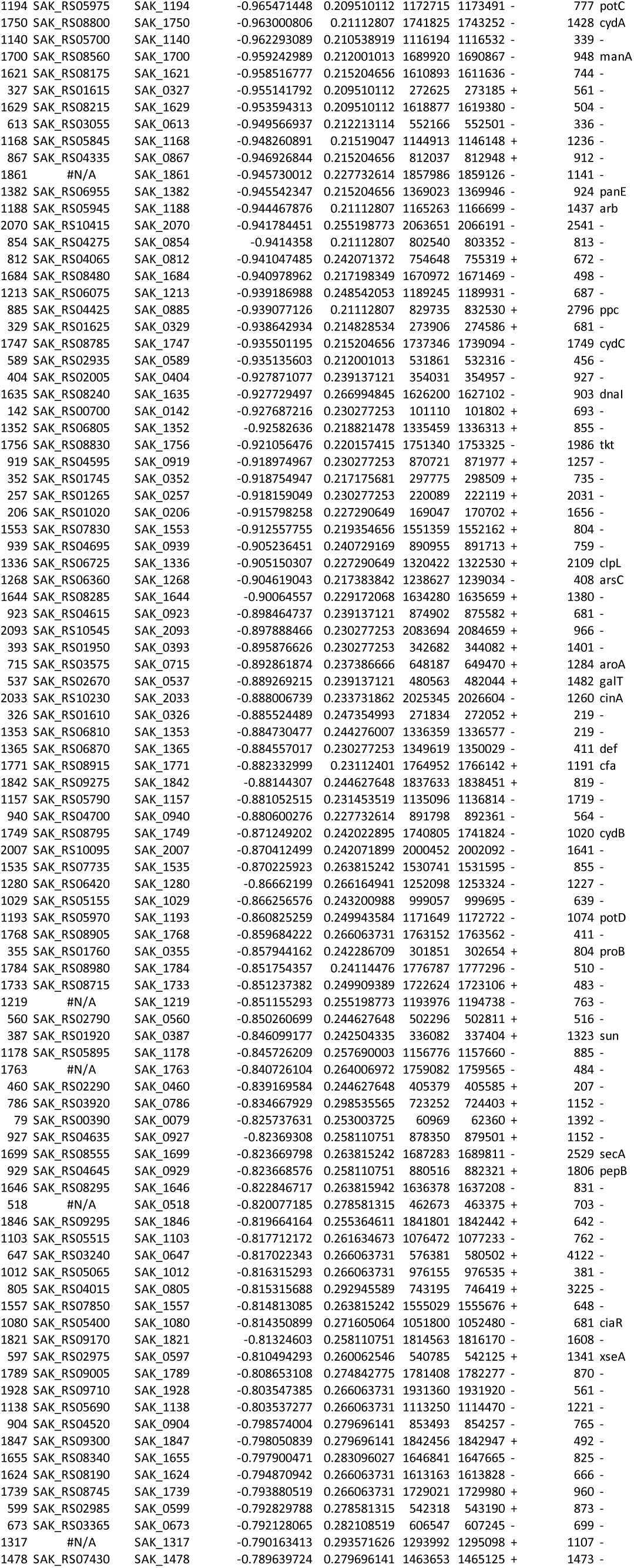
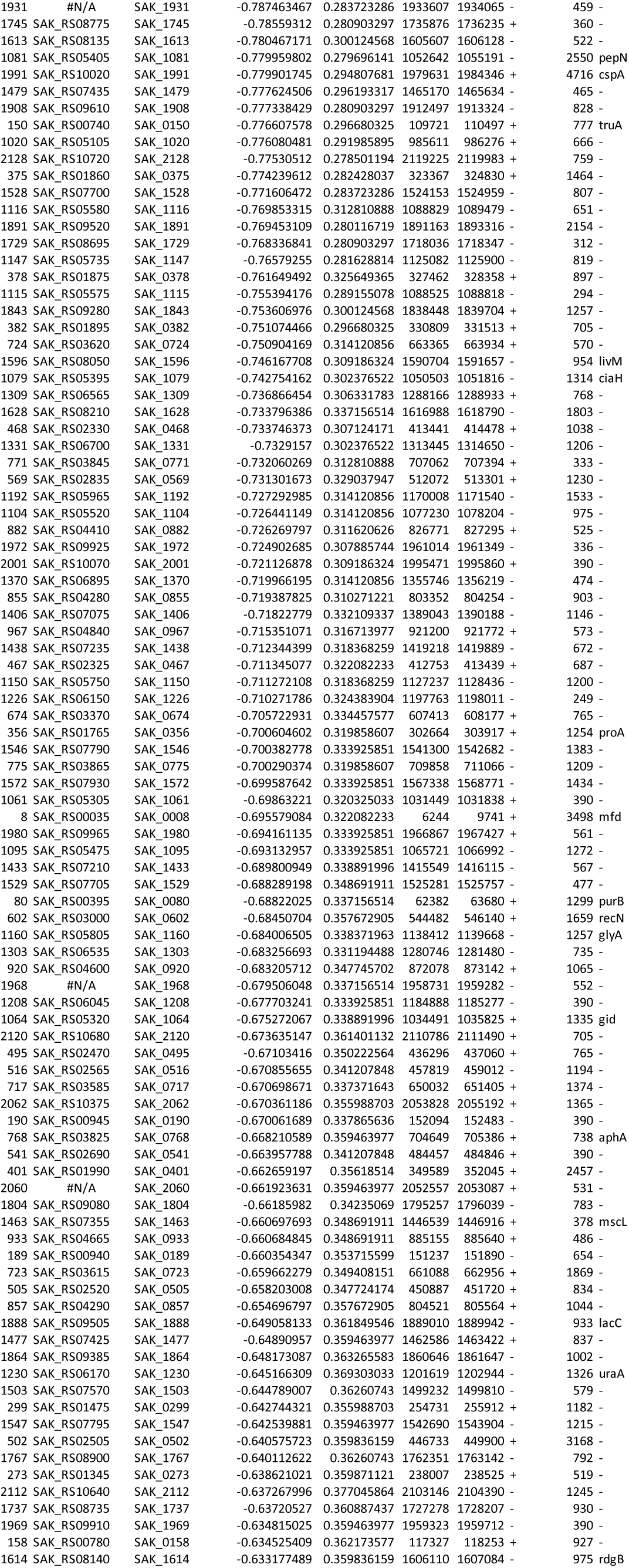
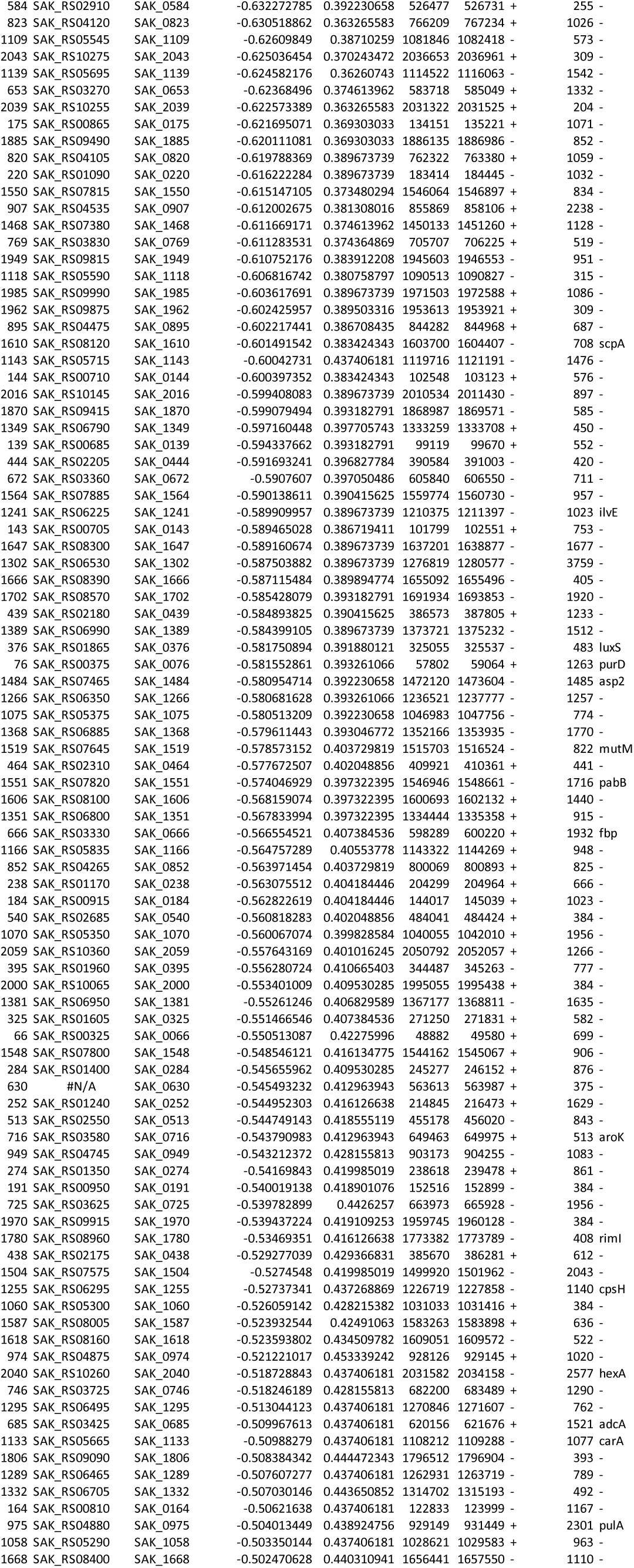
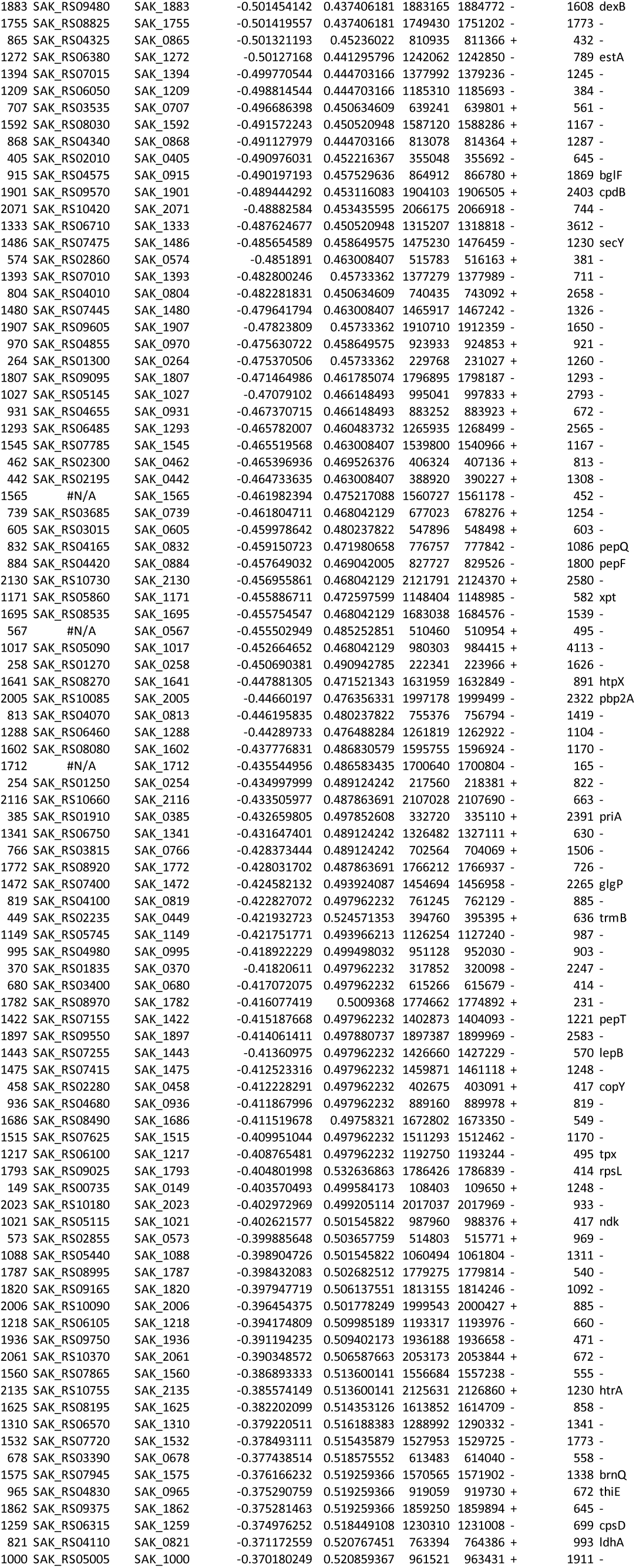
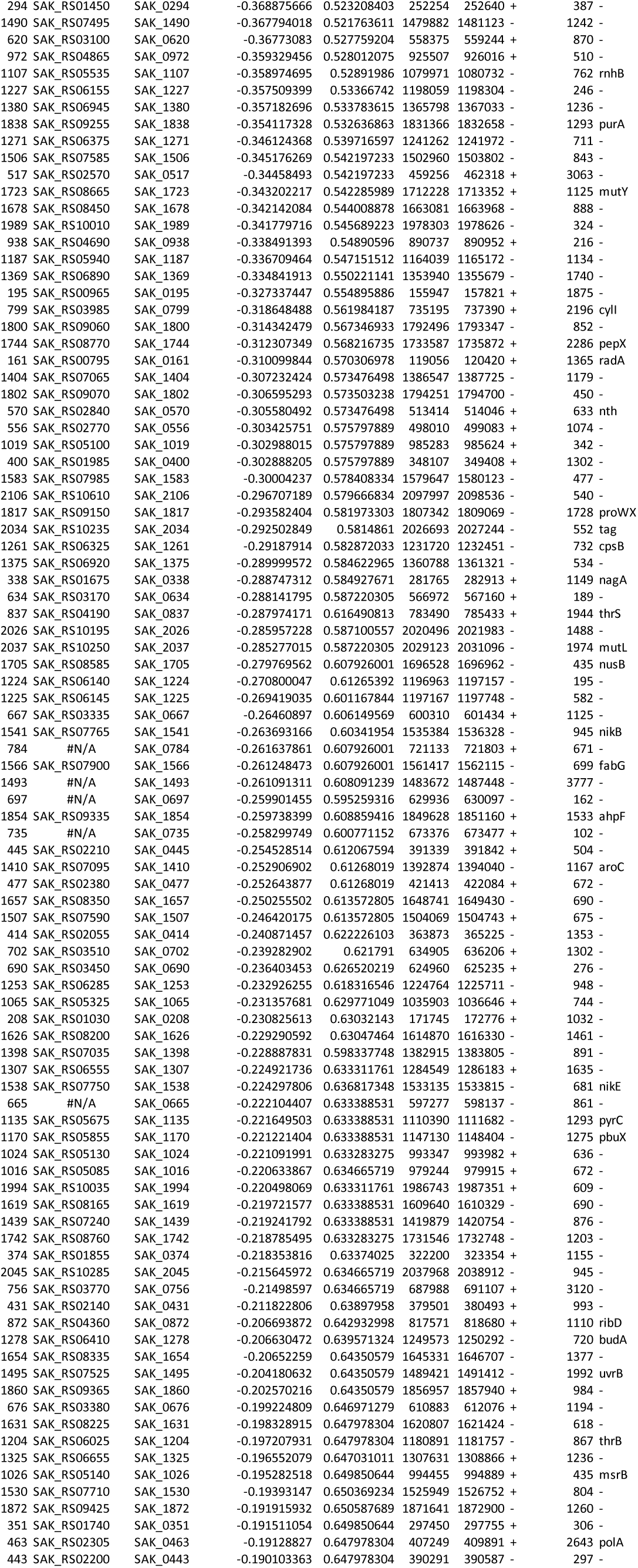
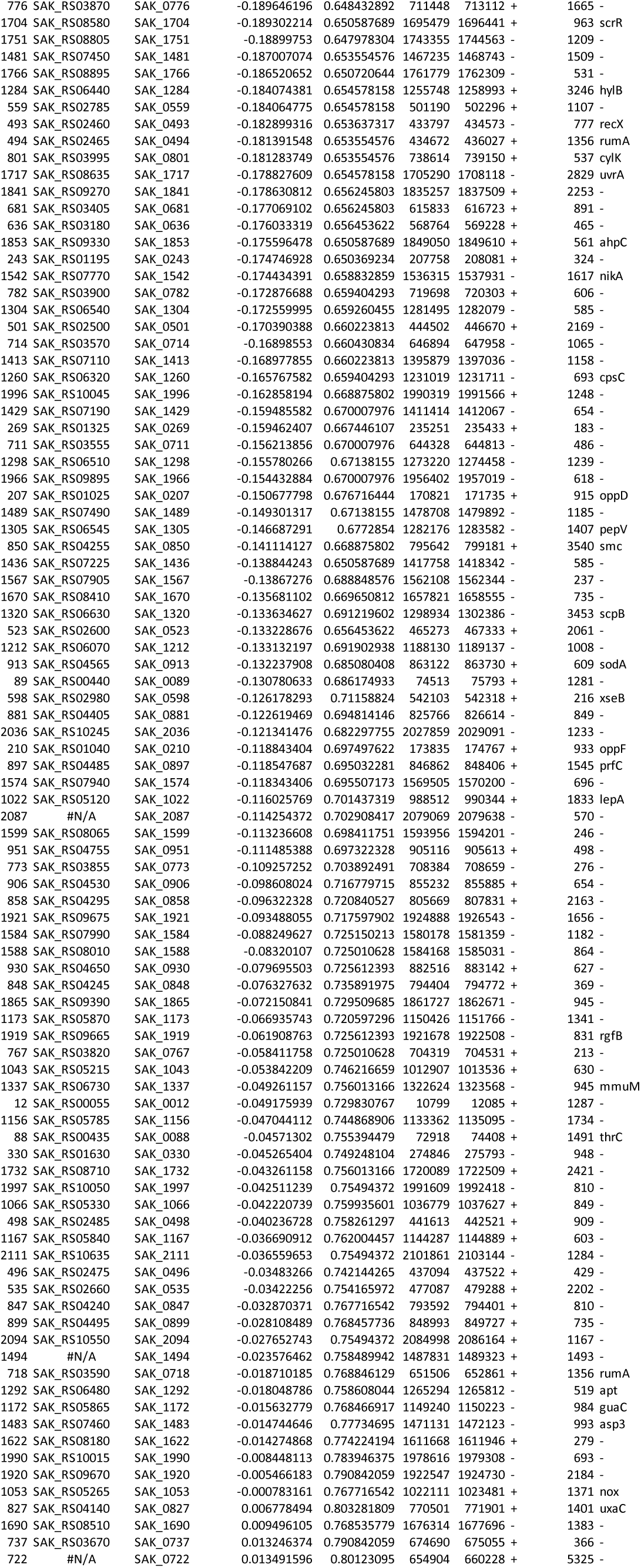
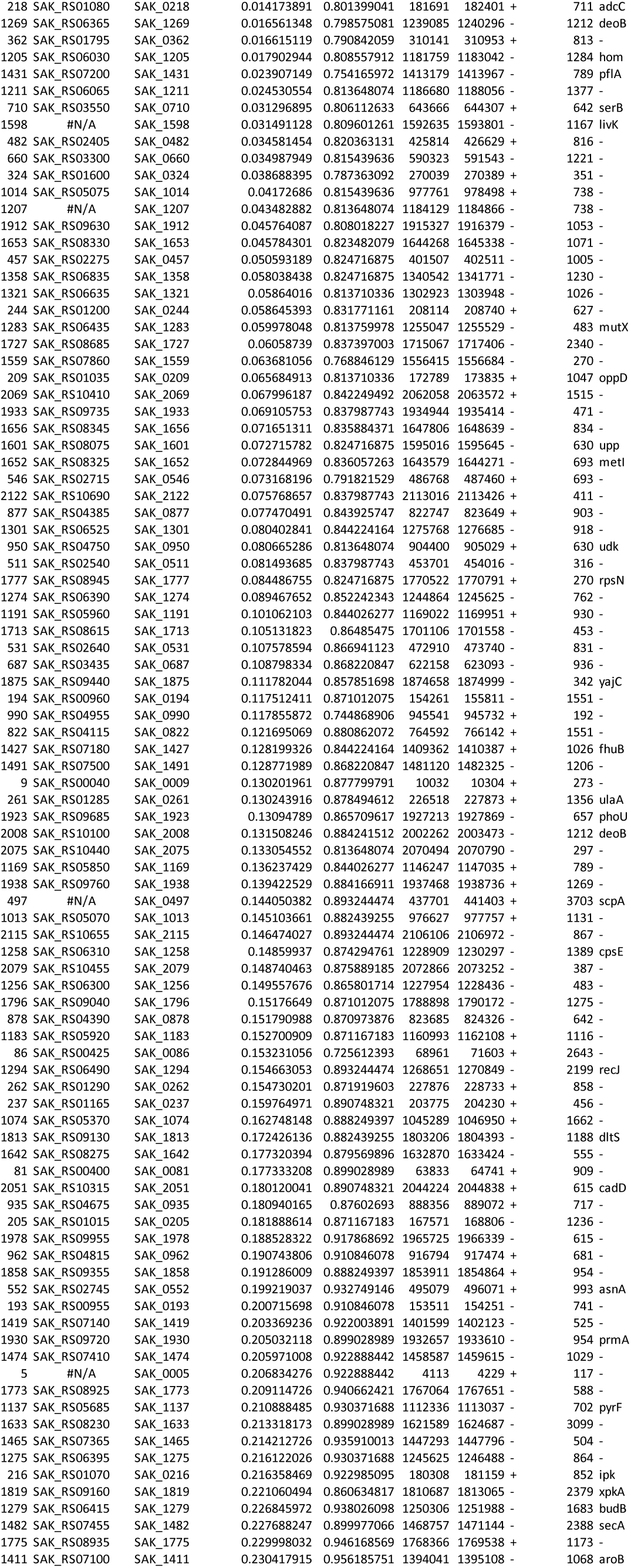
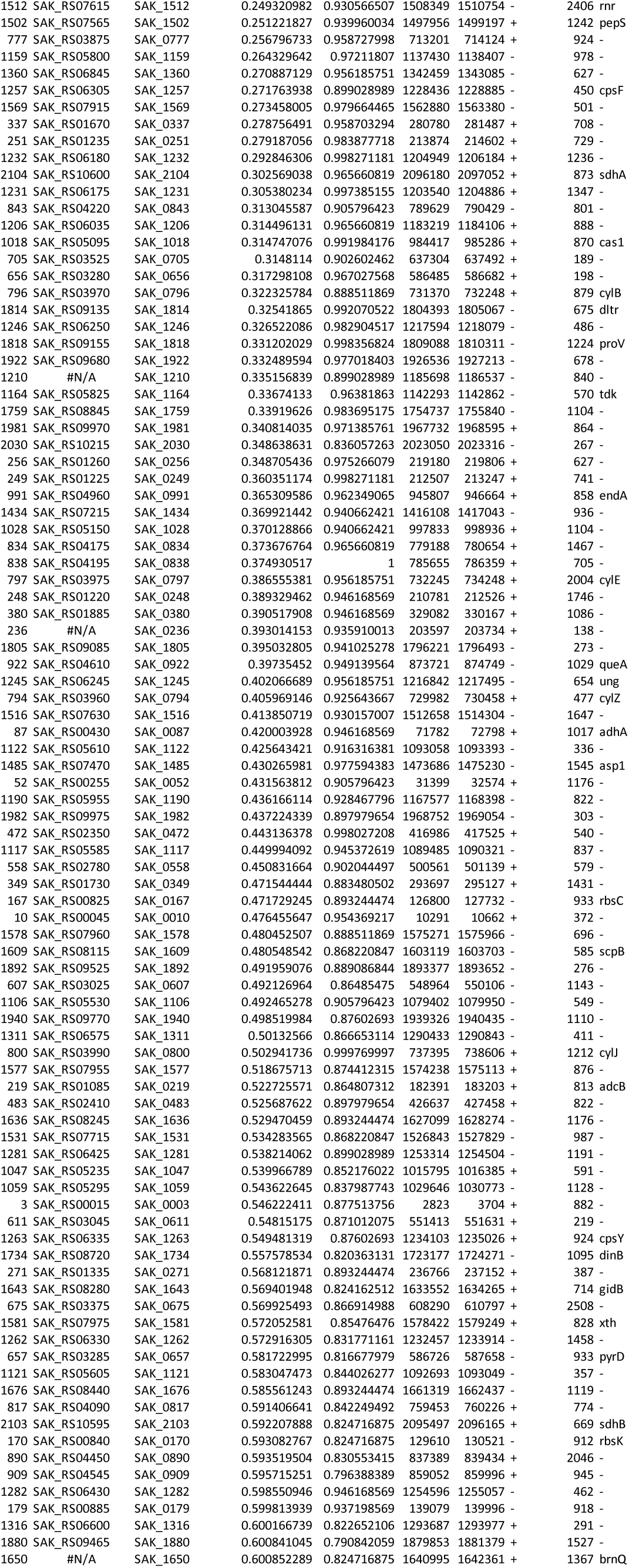
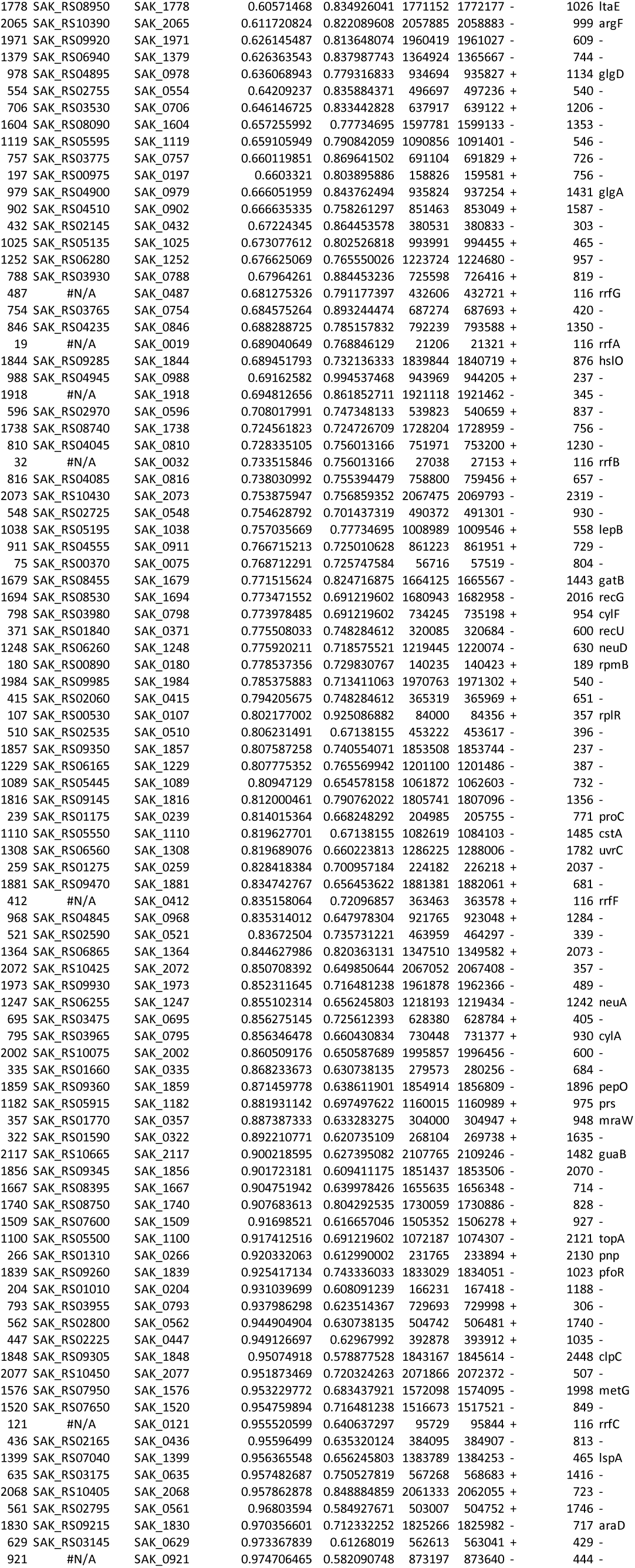
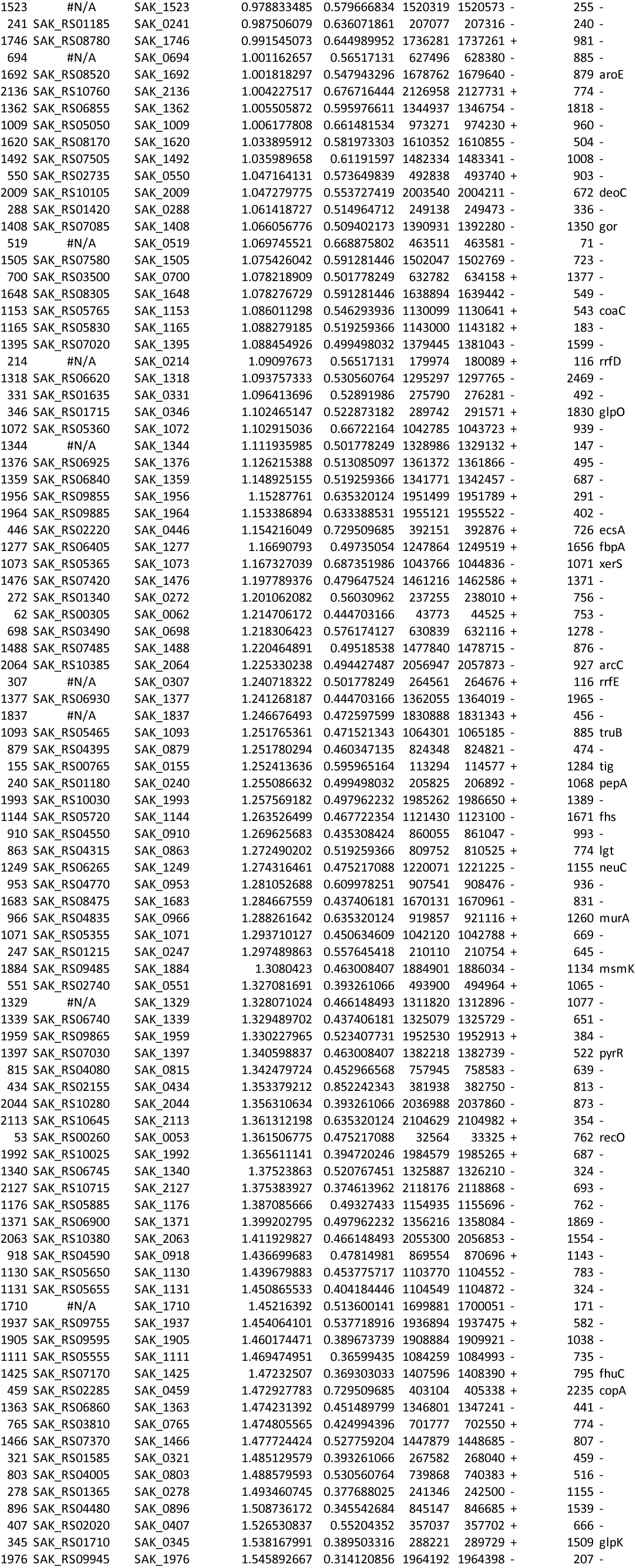
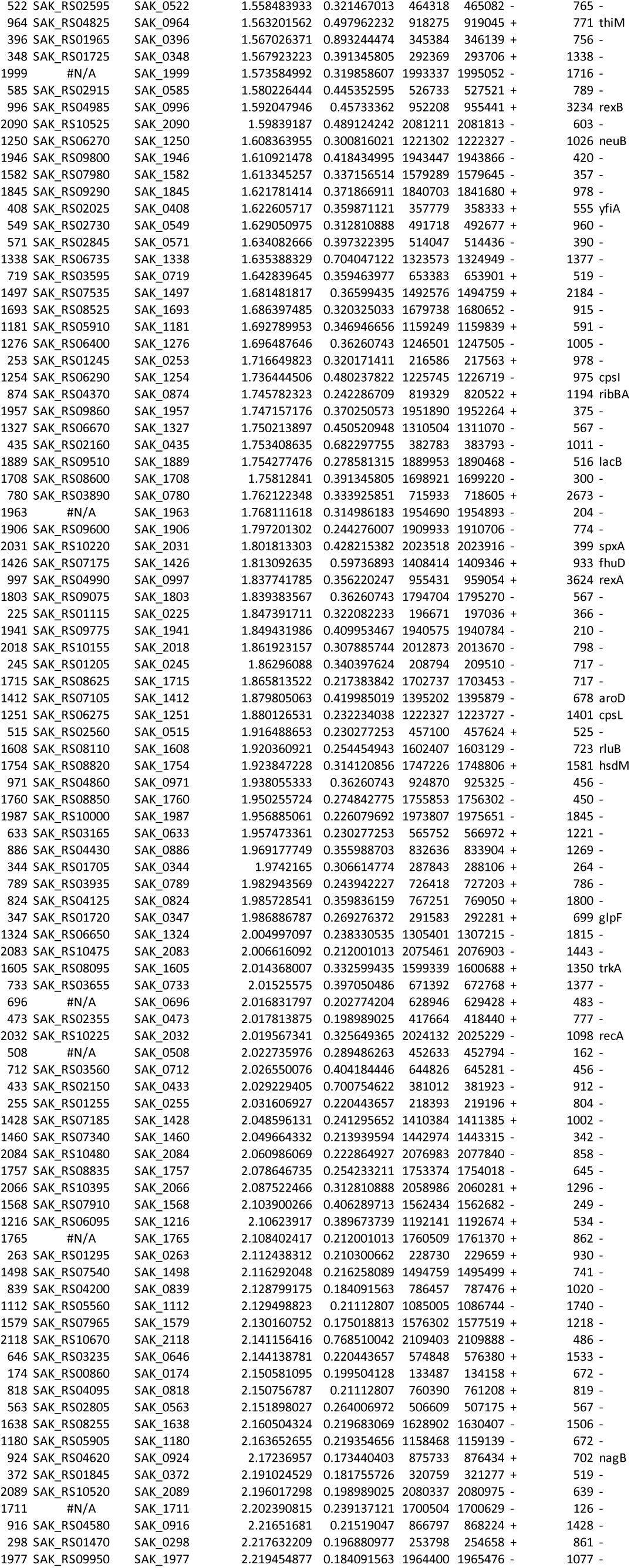
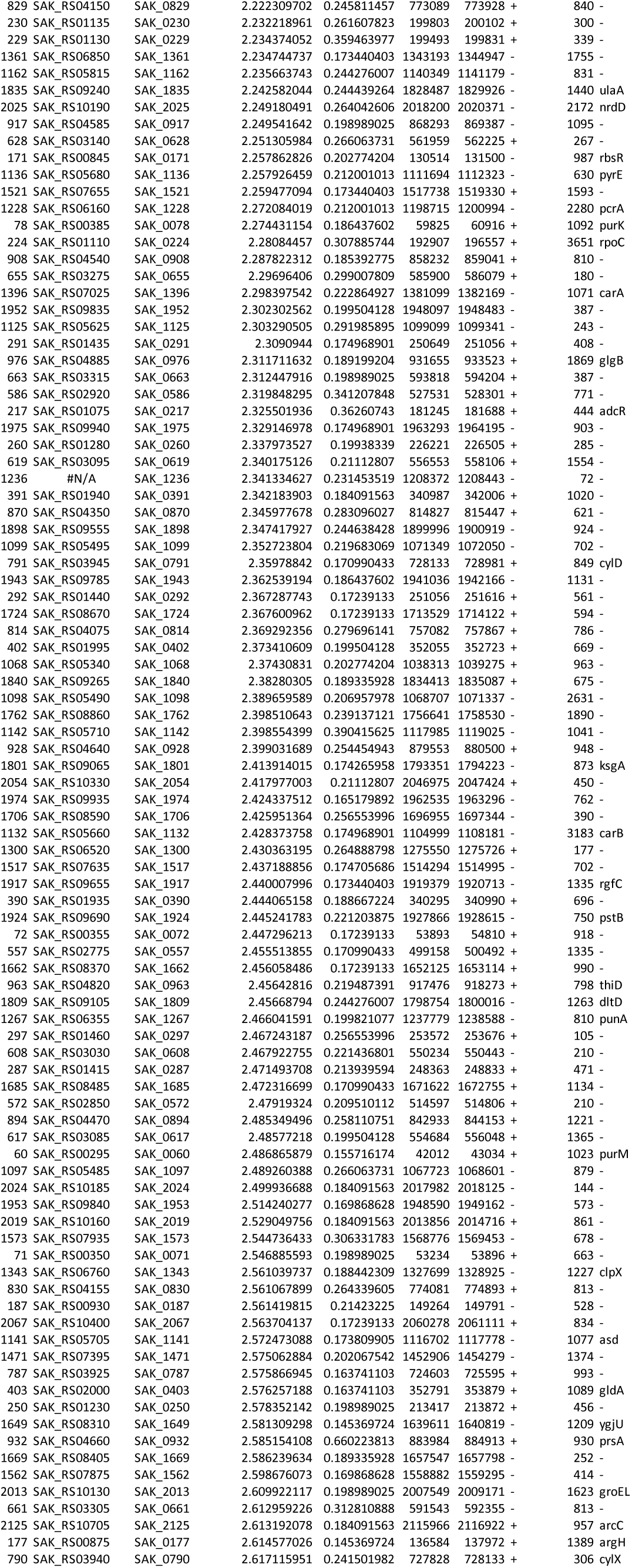
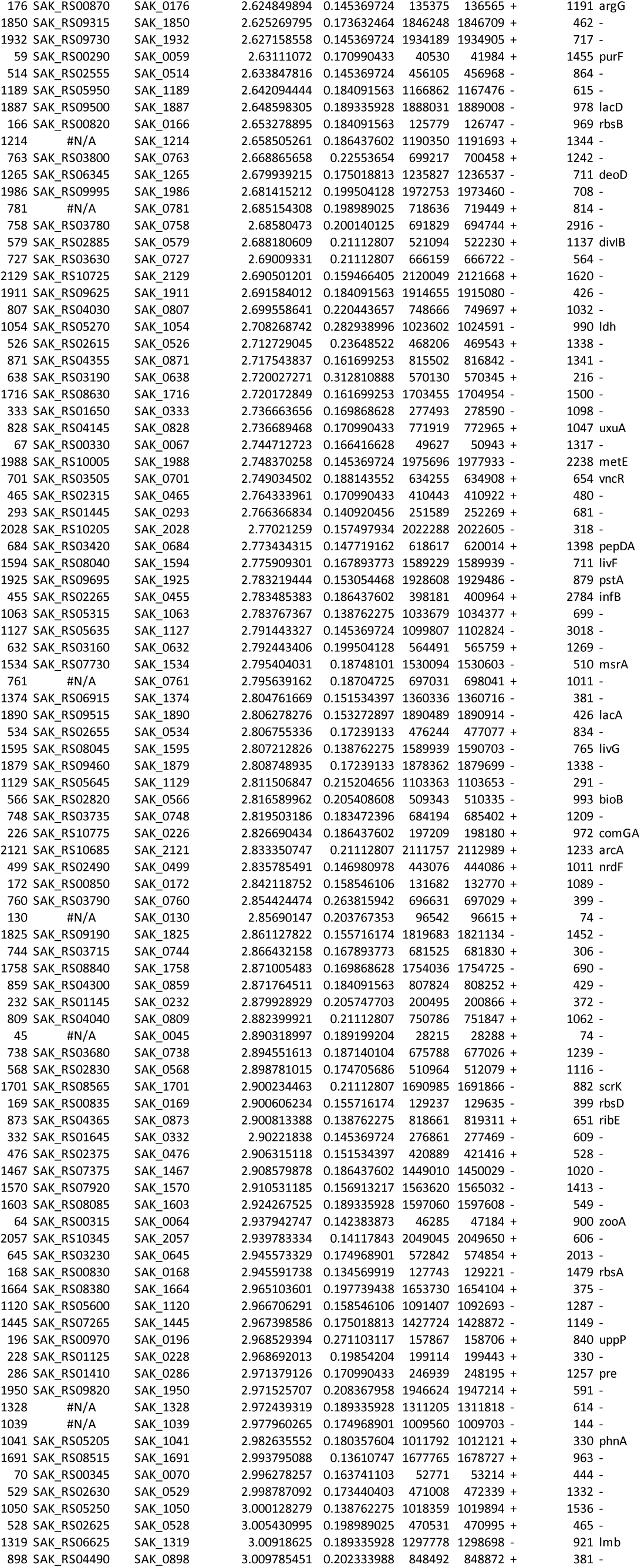
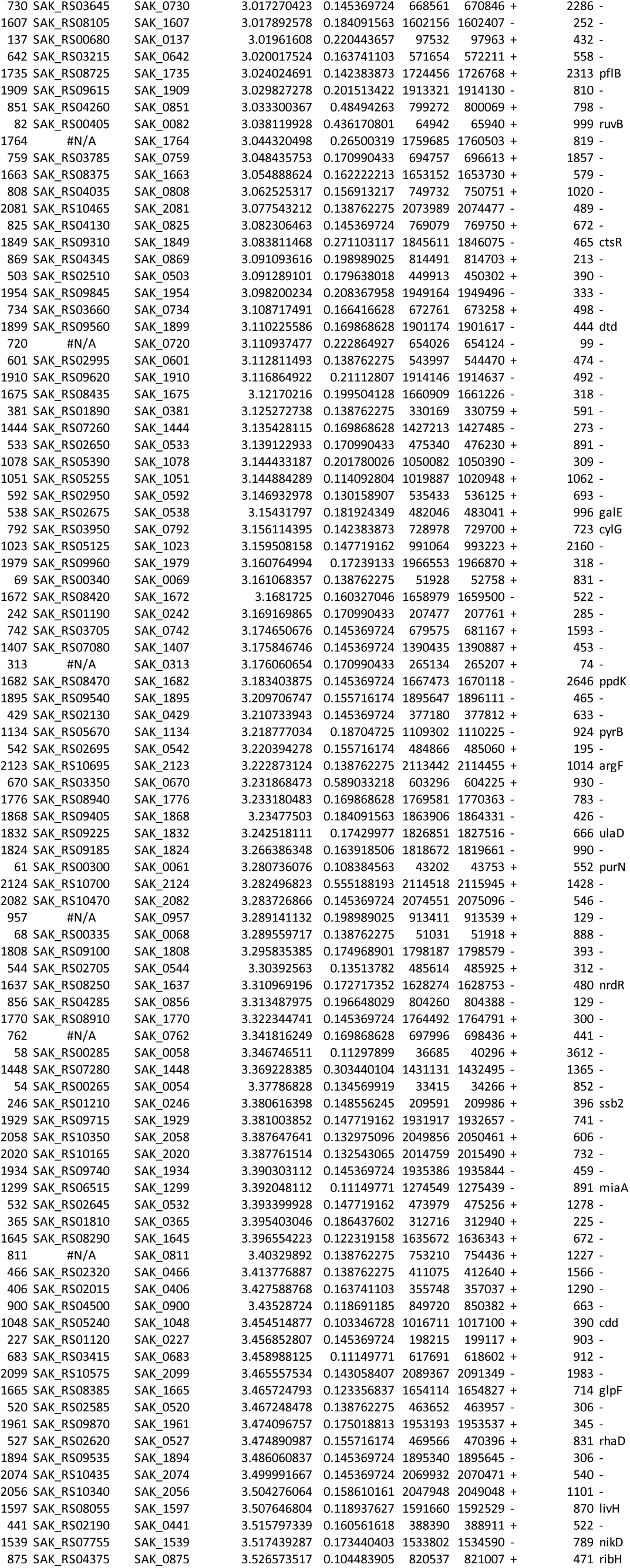
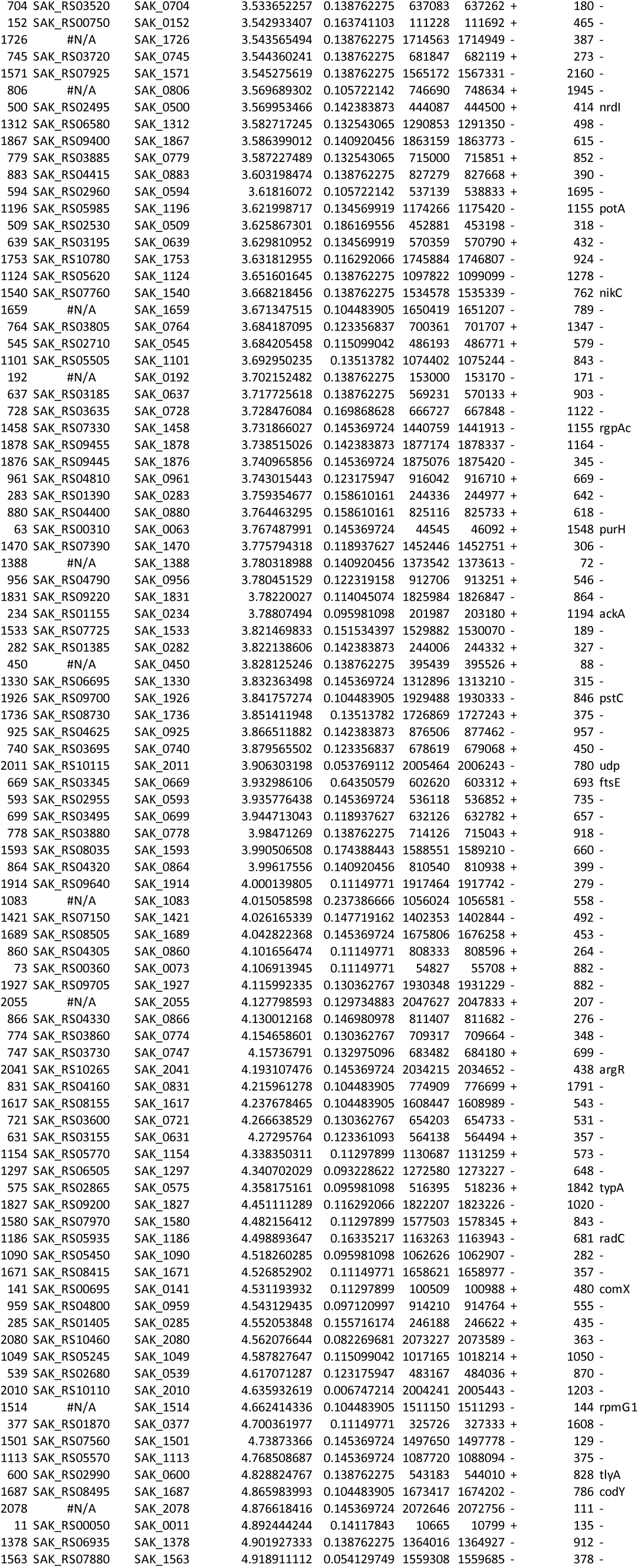
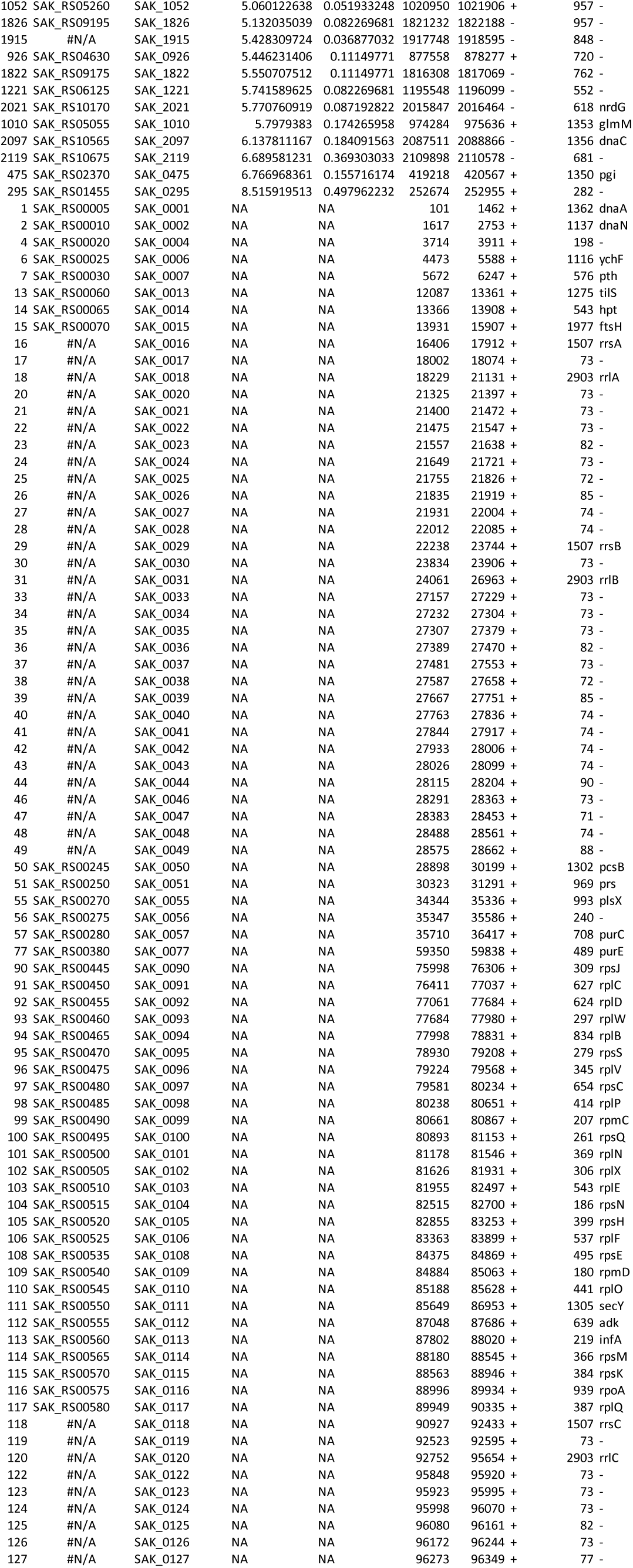
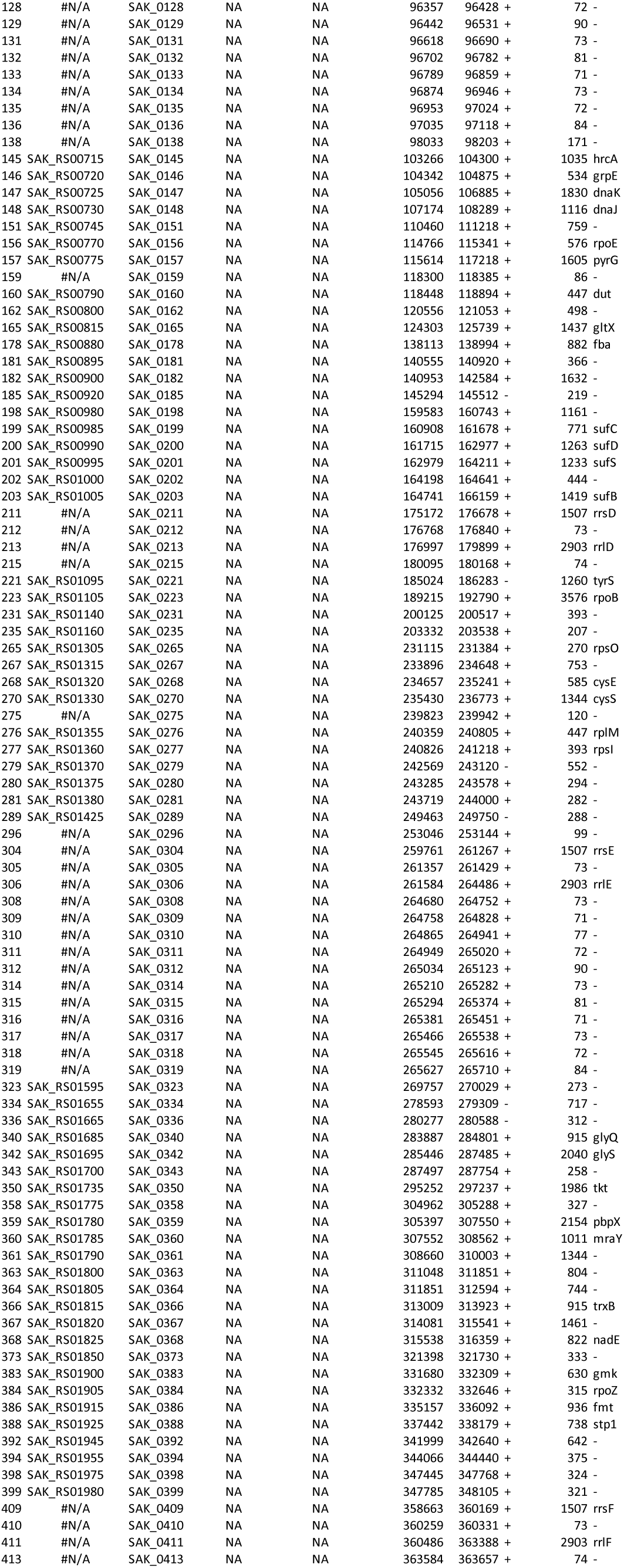
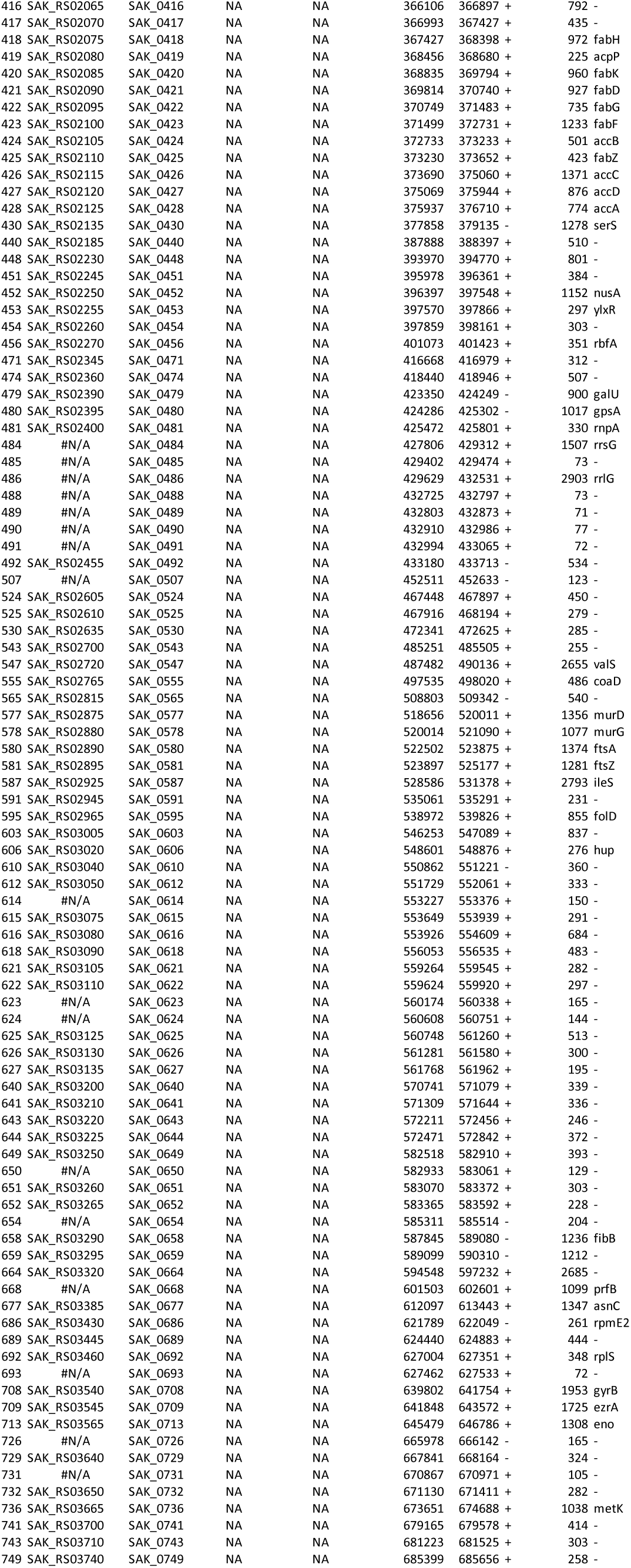
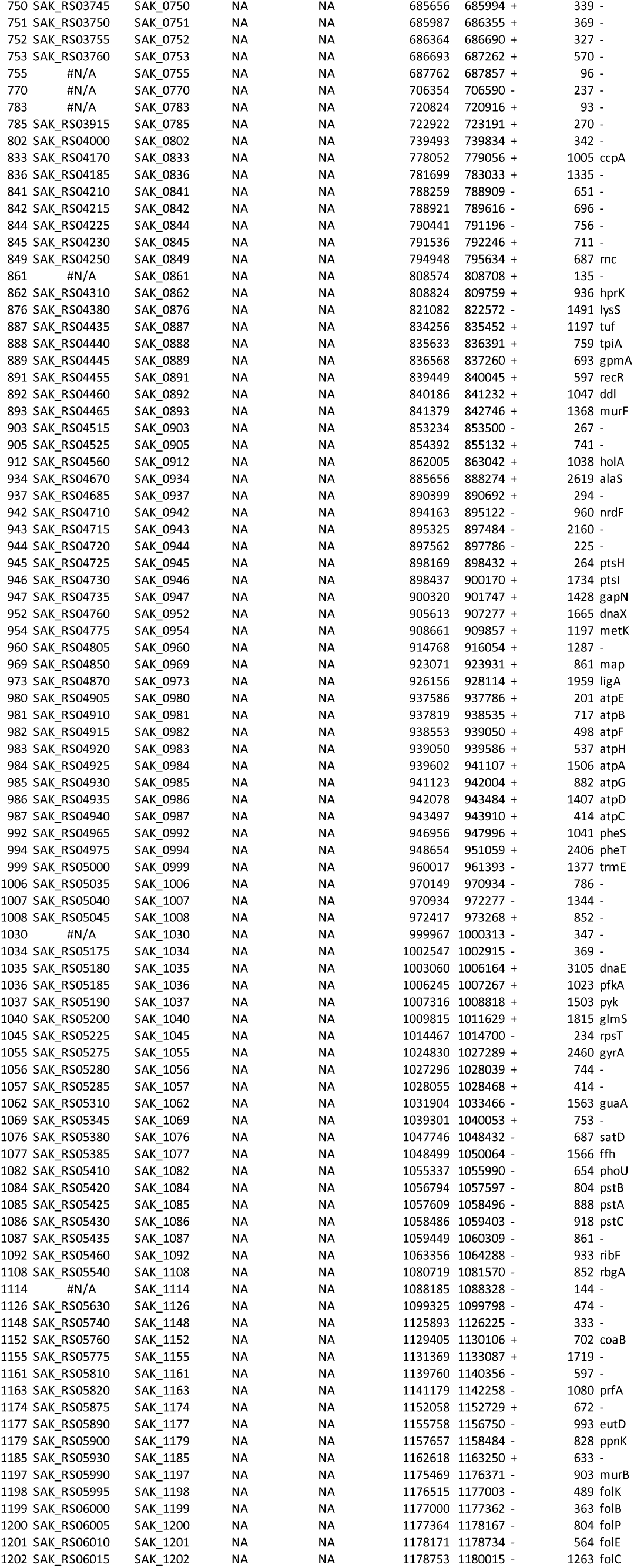
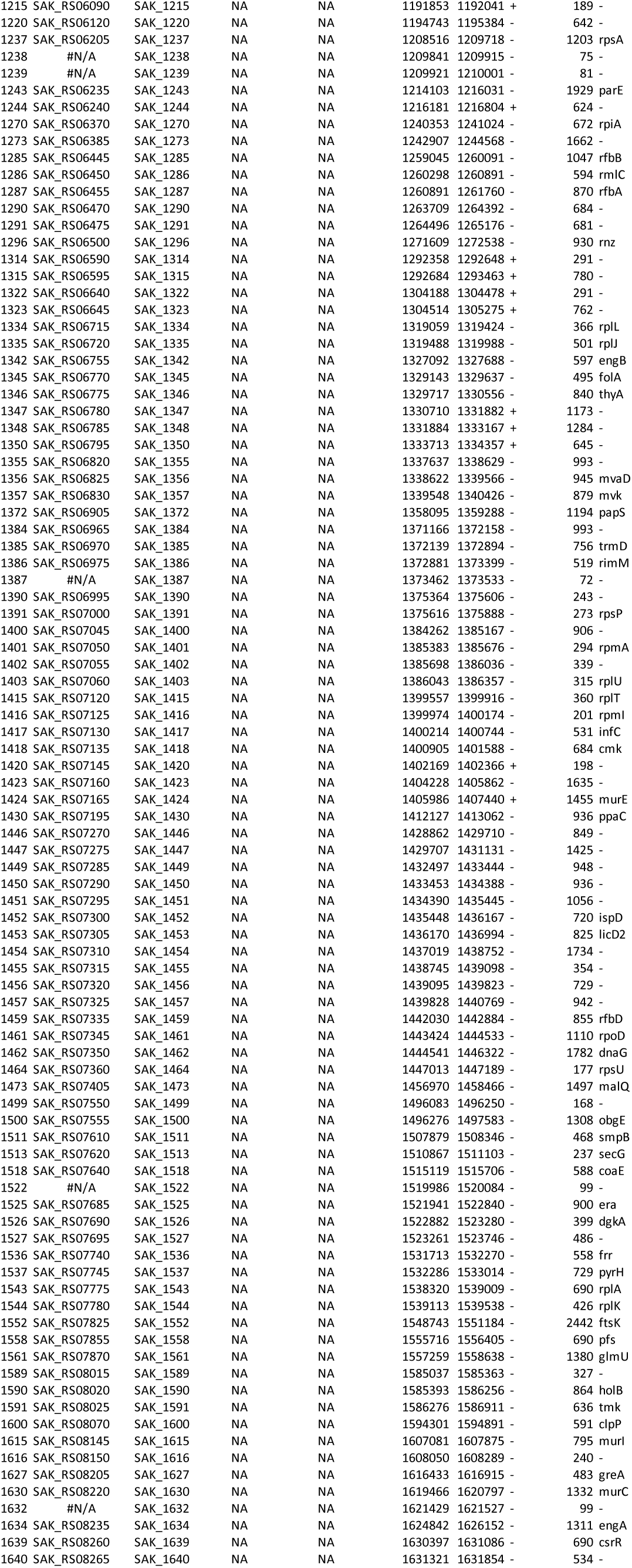
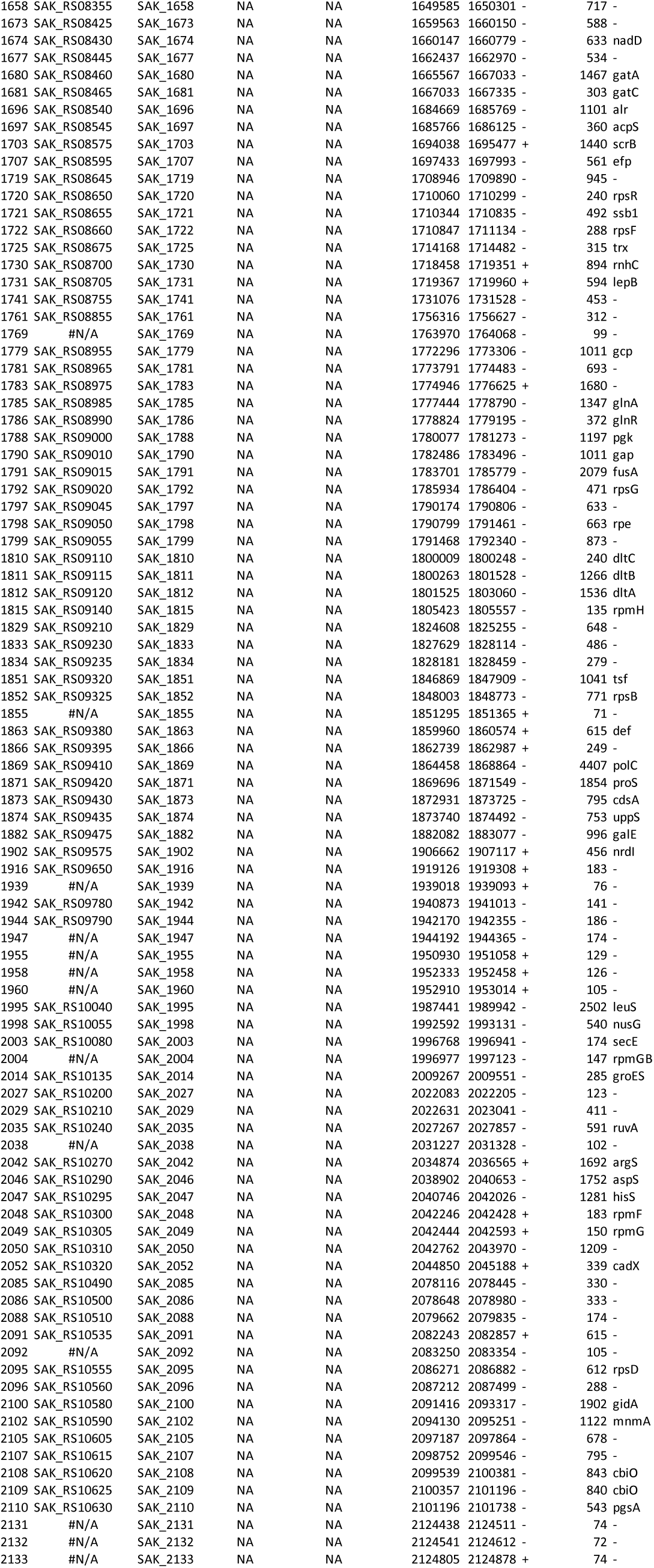

**Supplemental Data 2: RNA-seq comparing Δ*mrvR* with wild type 10/84 genome-wide gene expression** Output from DESeq analysis of RNA-seq read alignments. Log_2_ fold-change and adjusted p values are provided for three growth phases as described in the text.

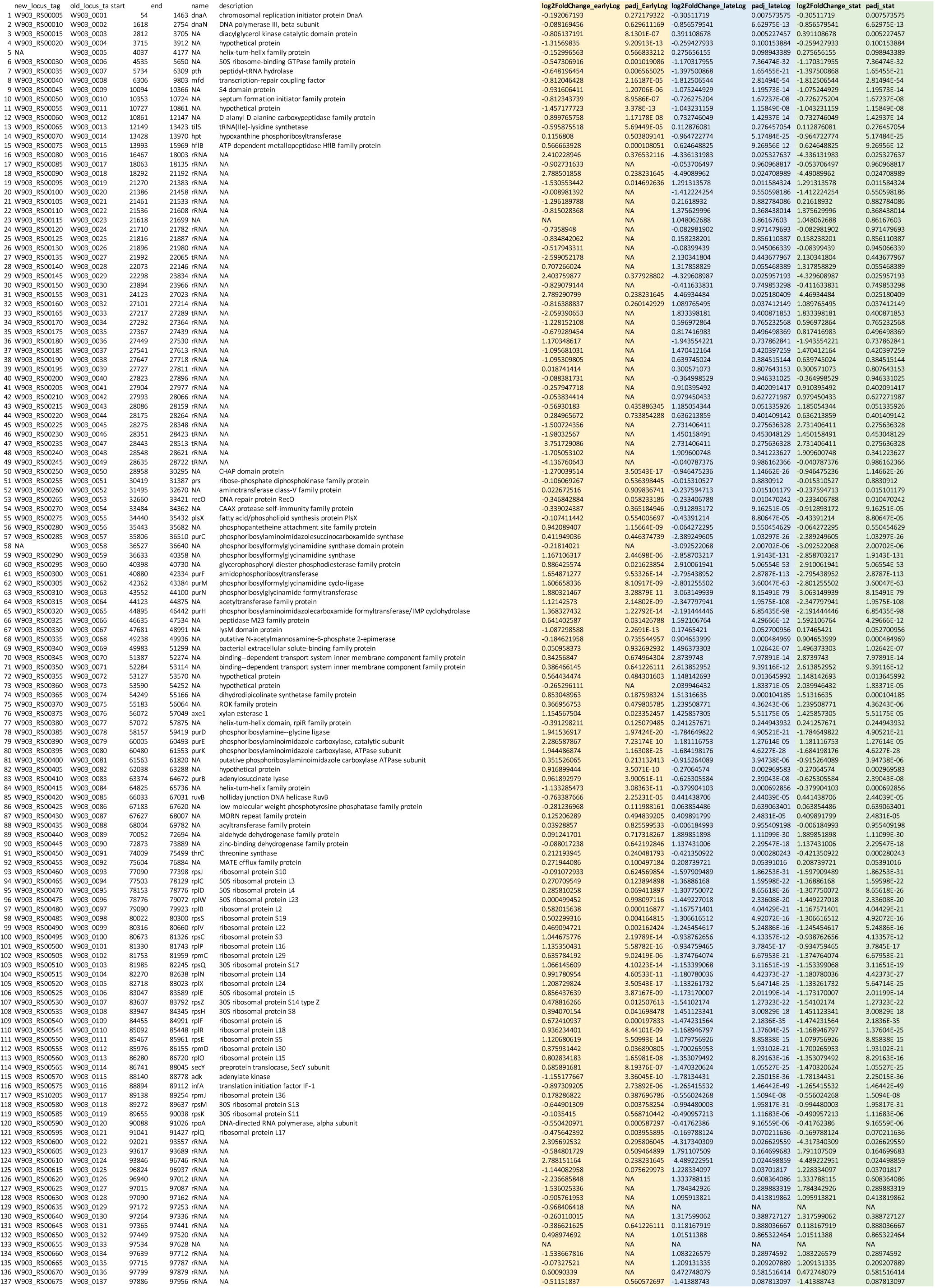
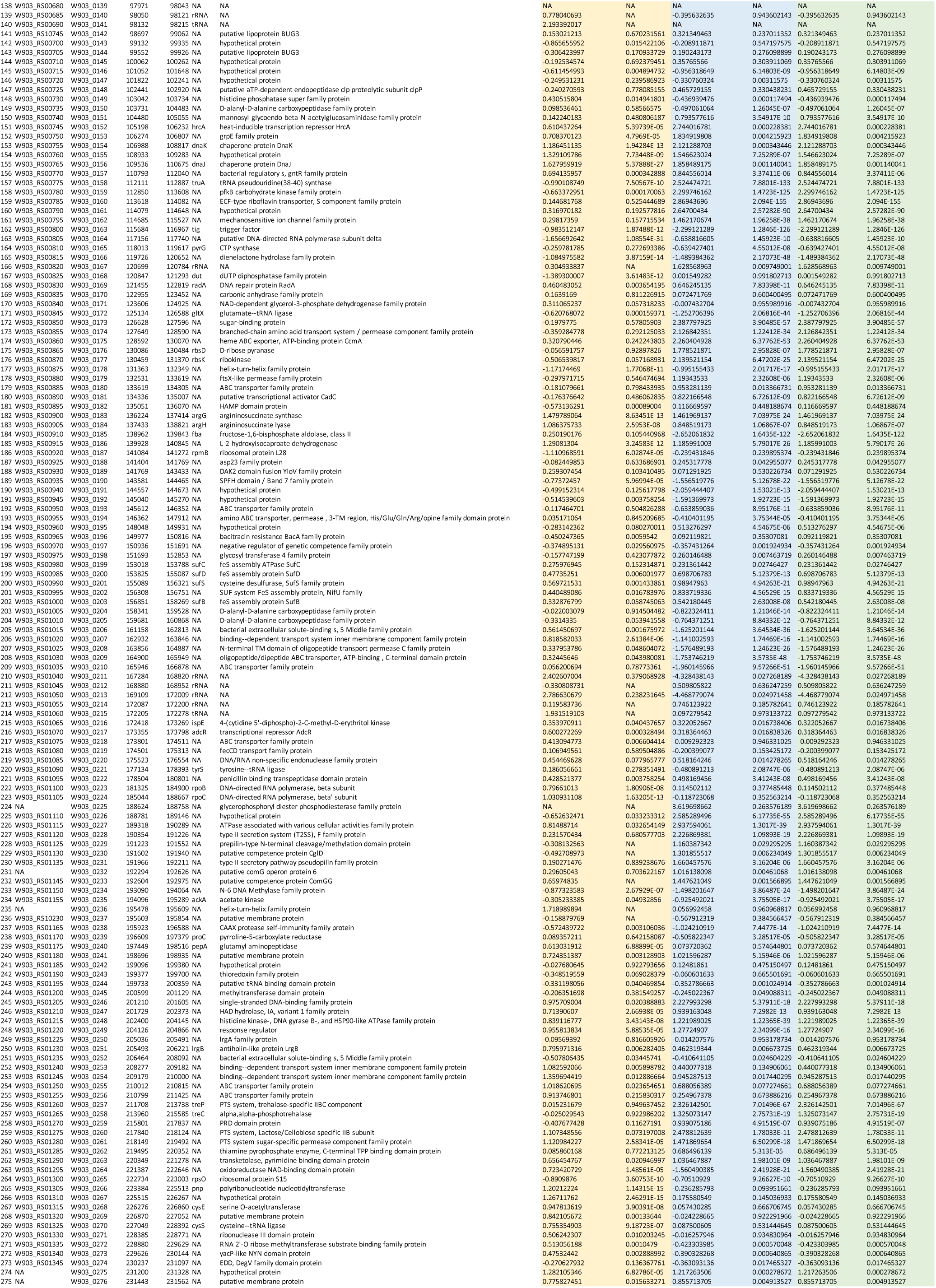
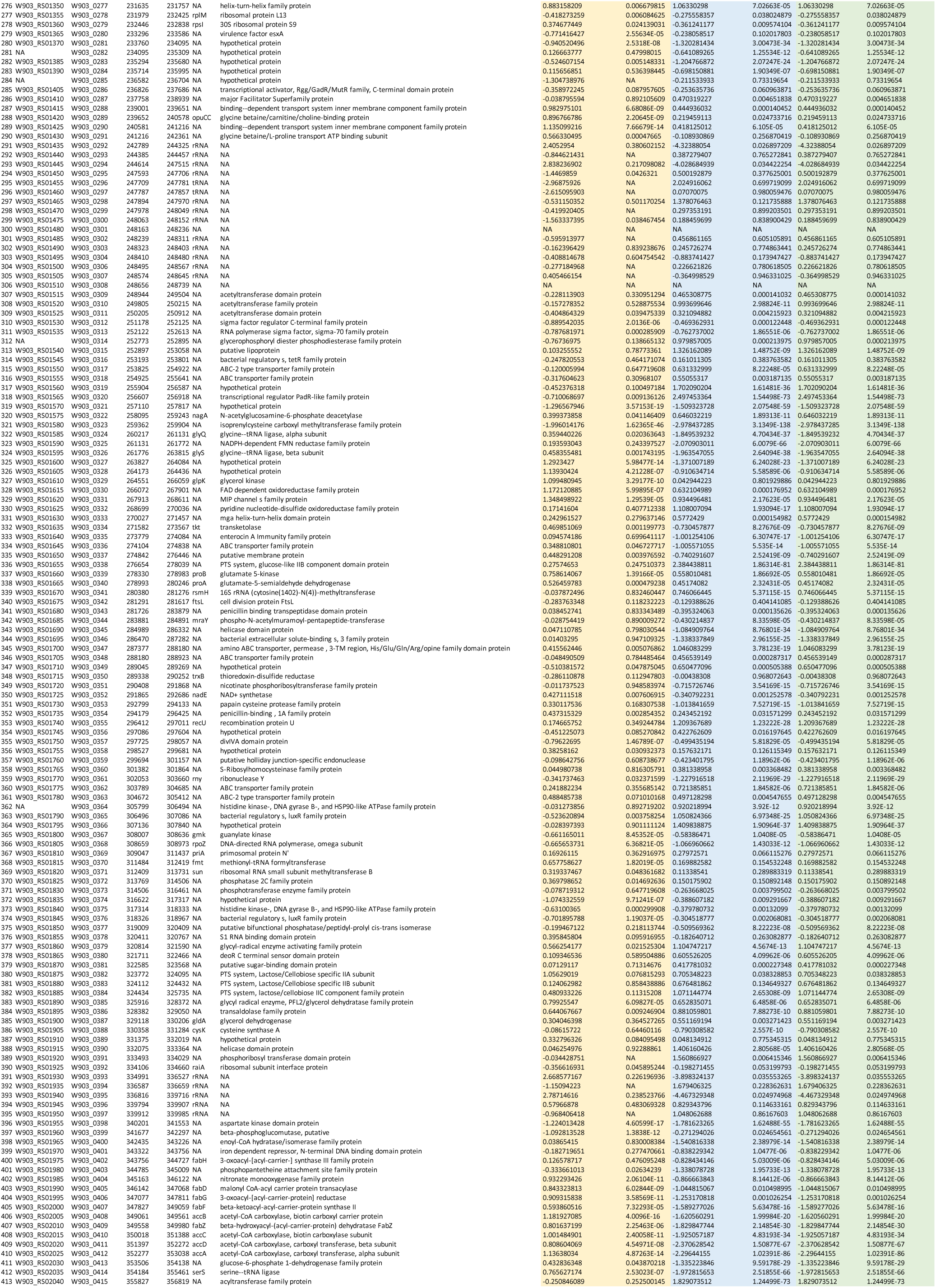
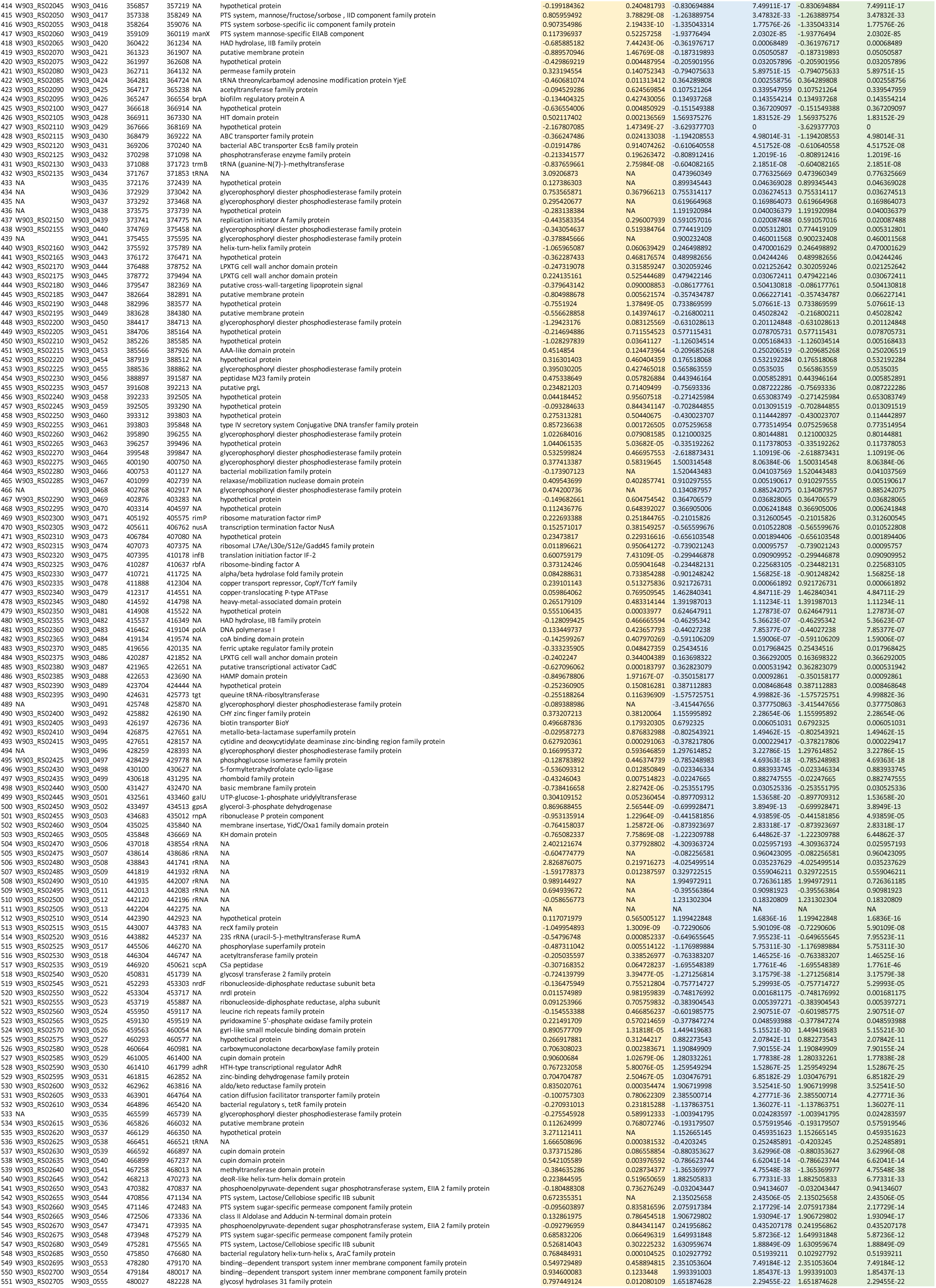
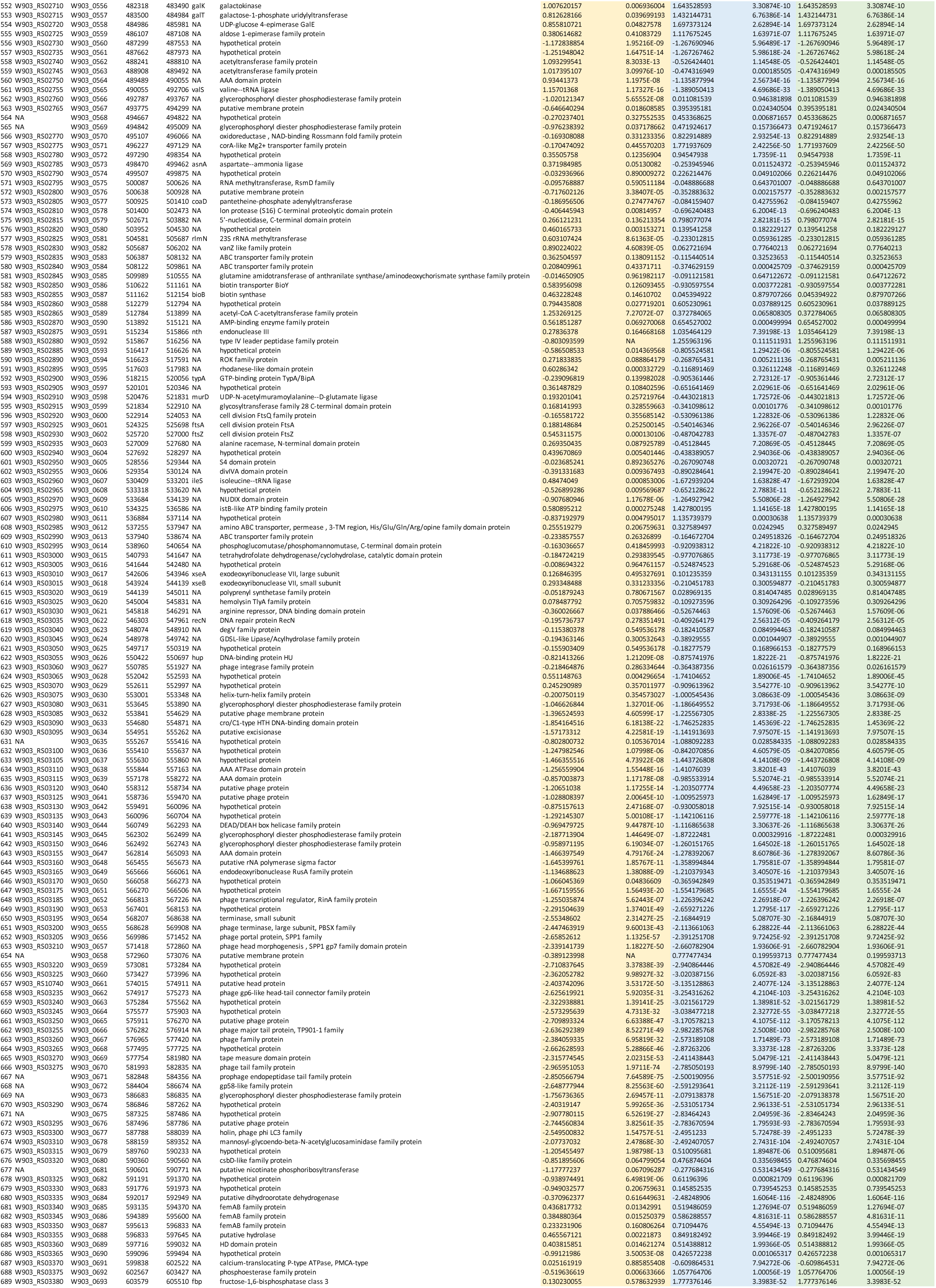
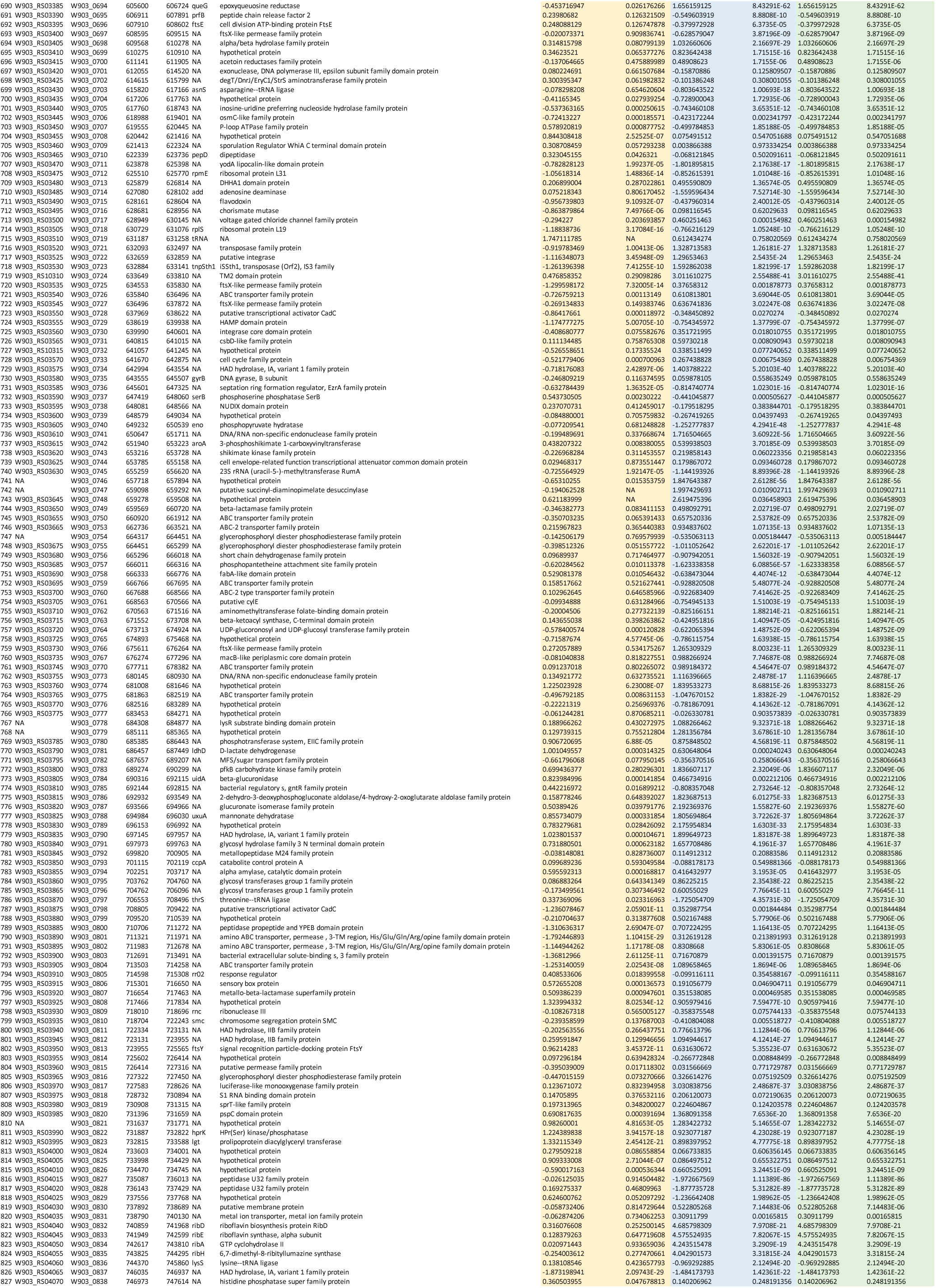
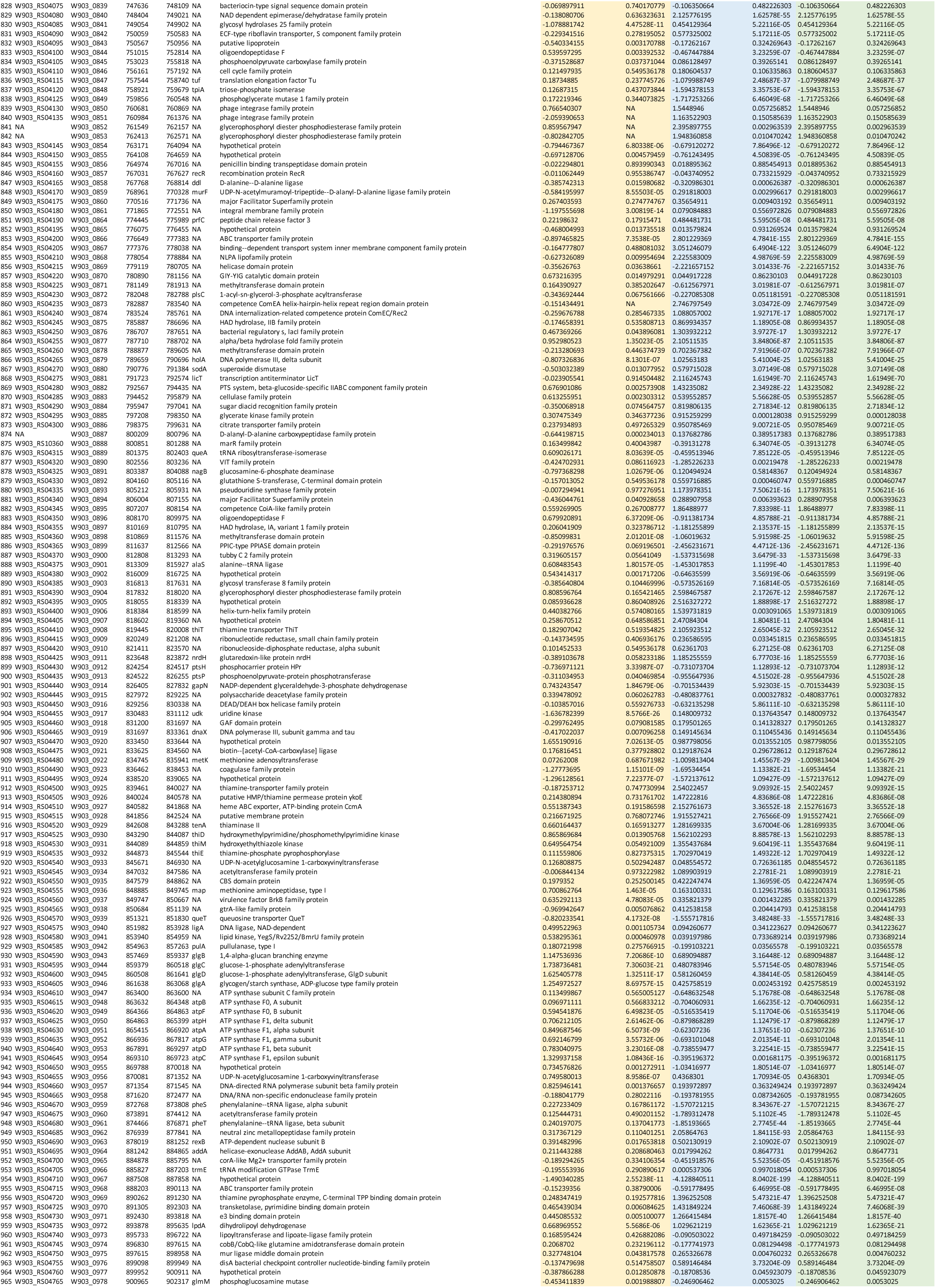
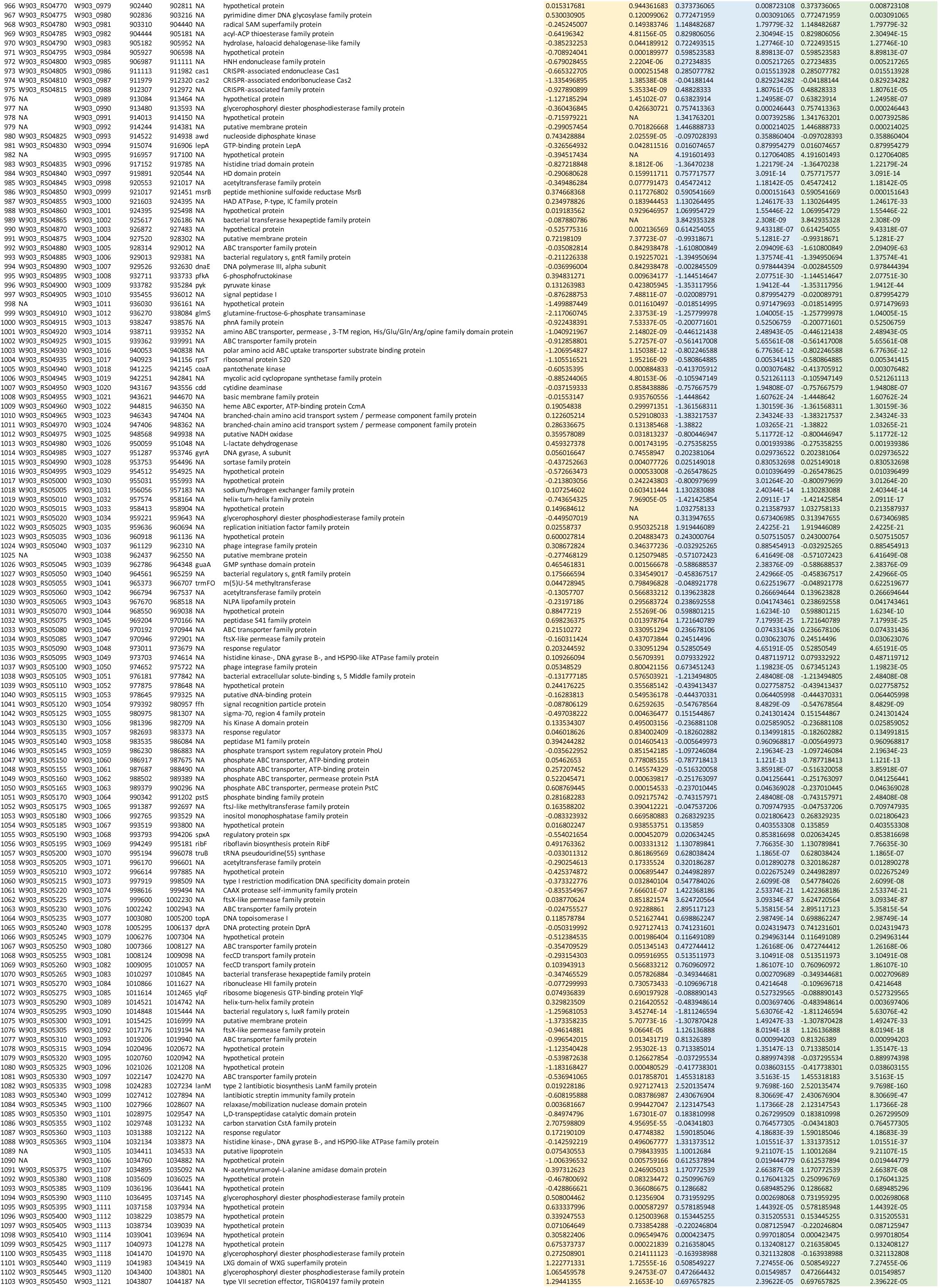
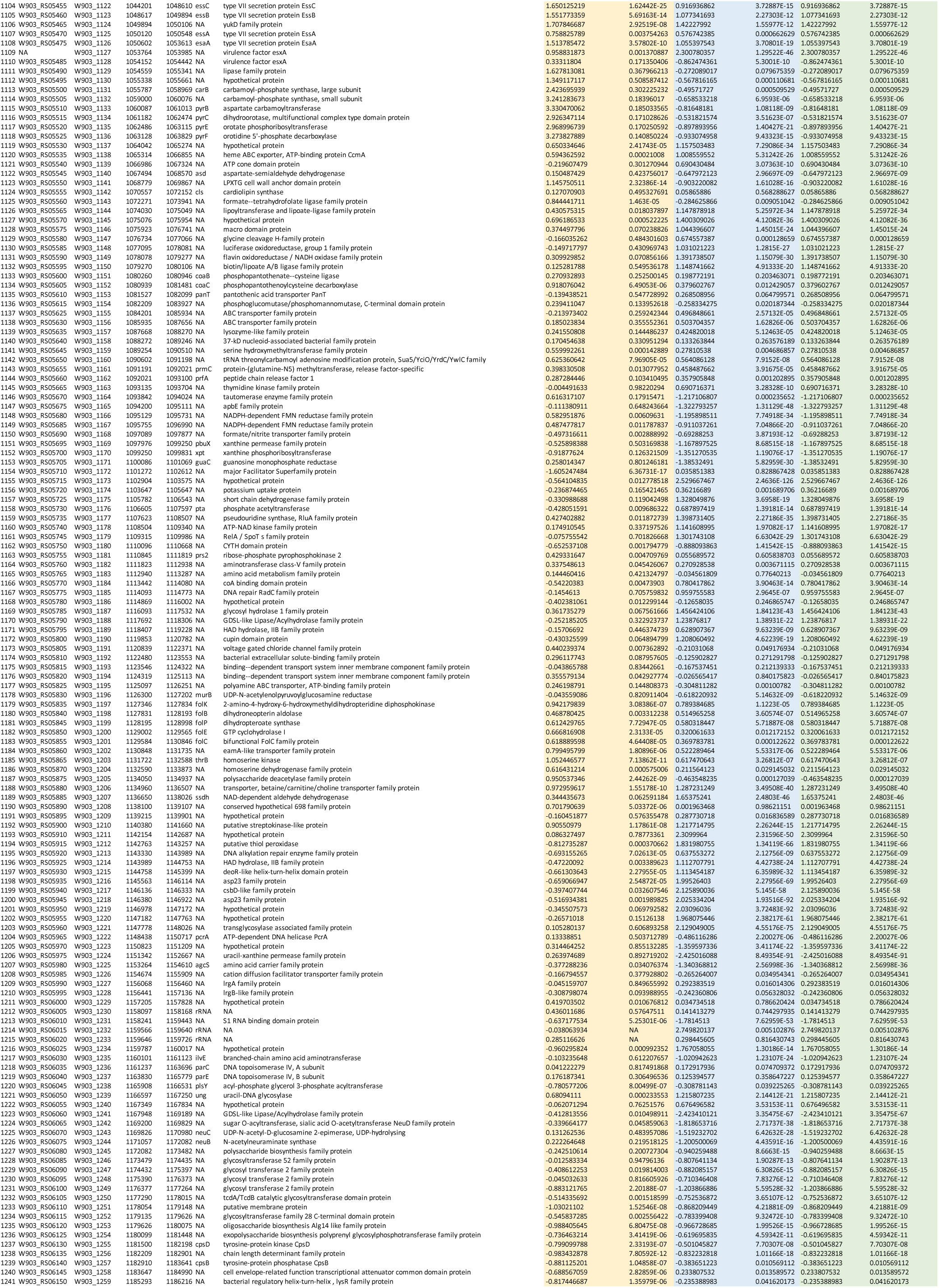
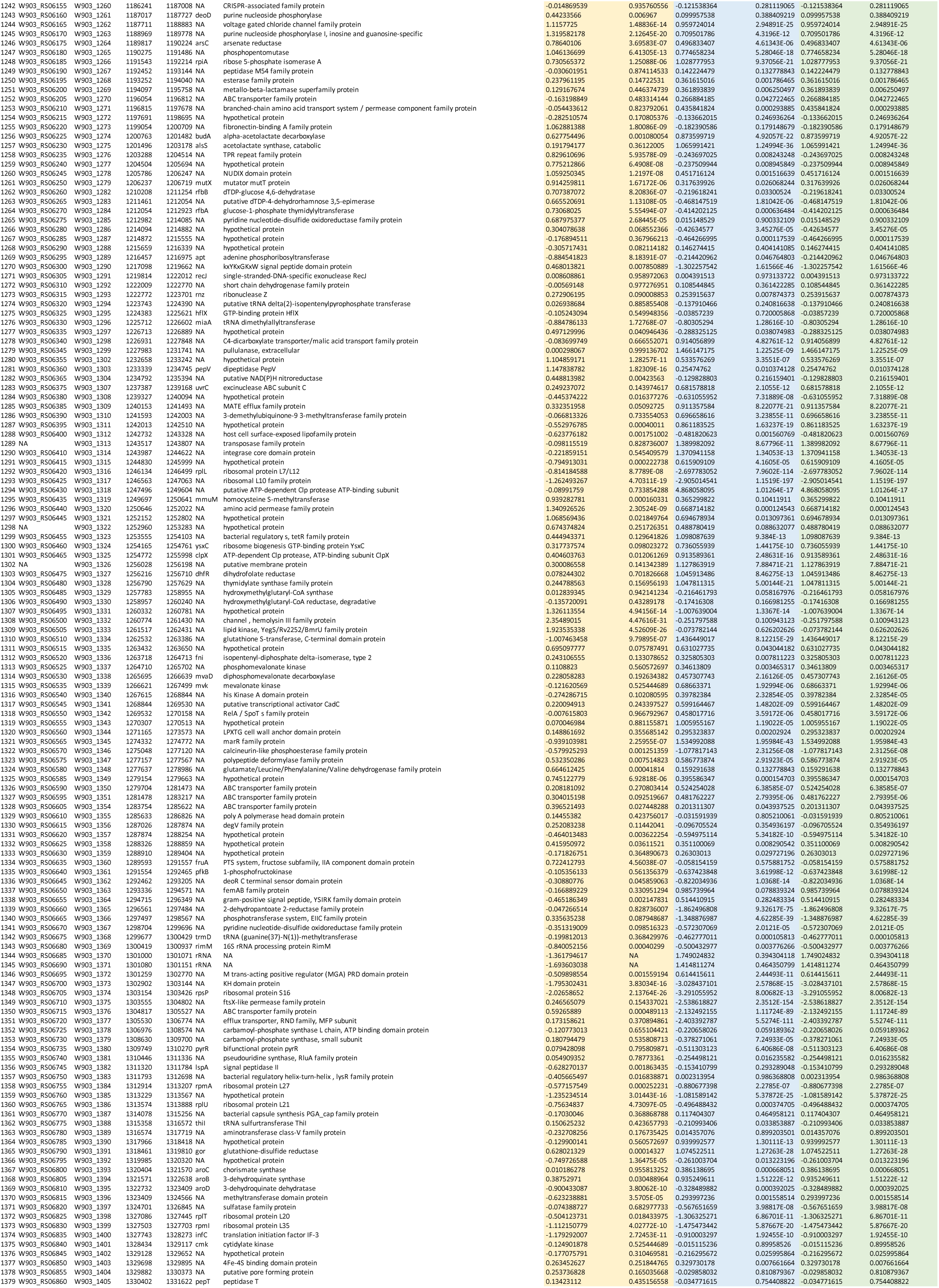
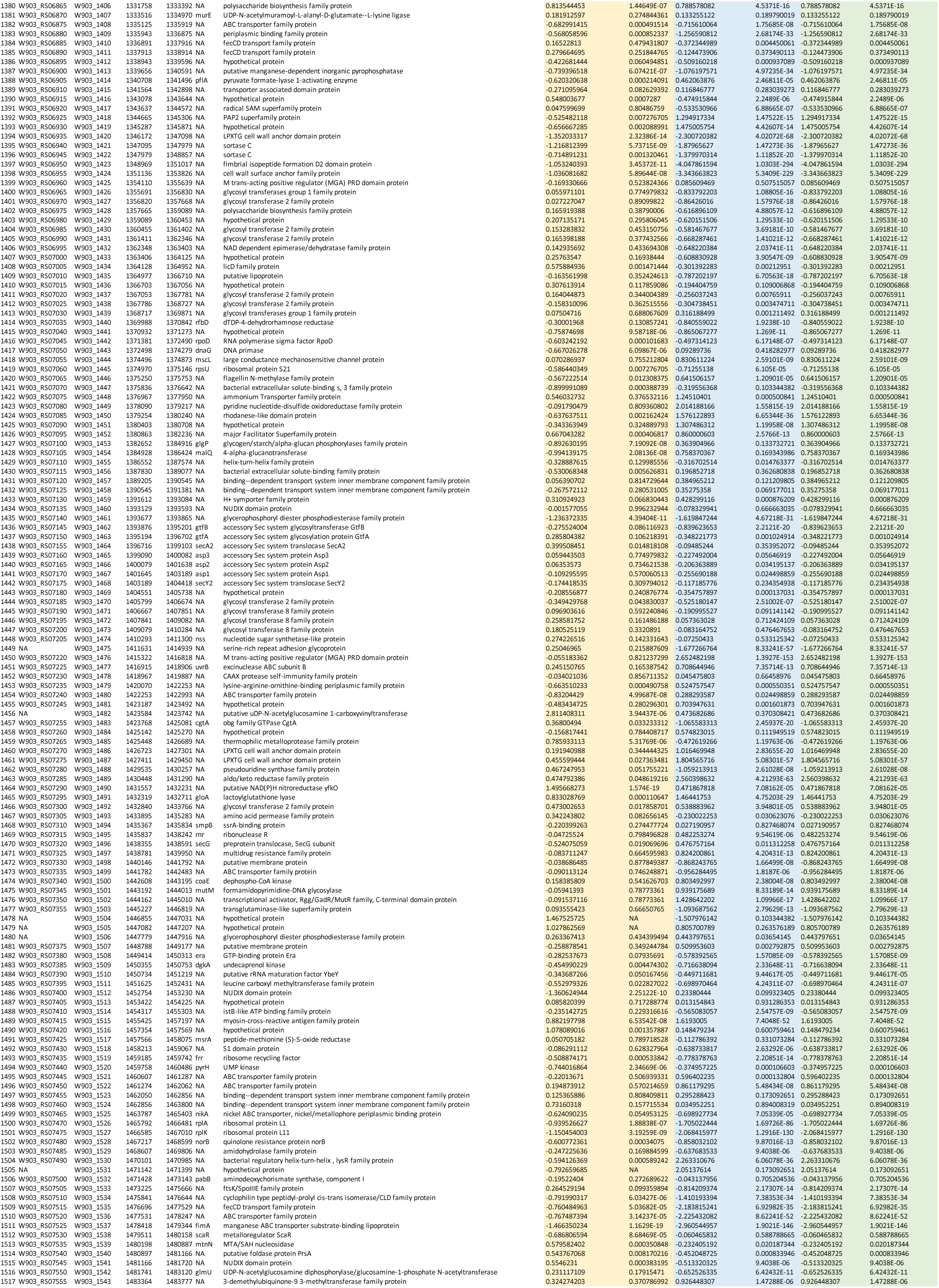
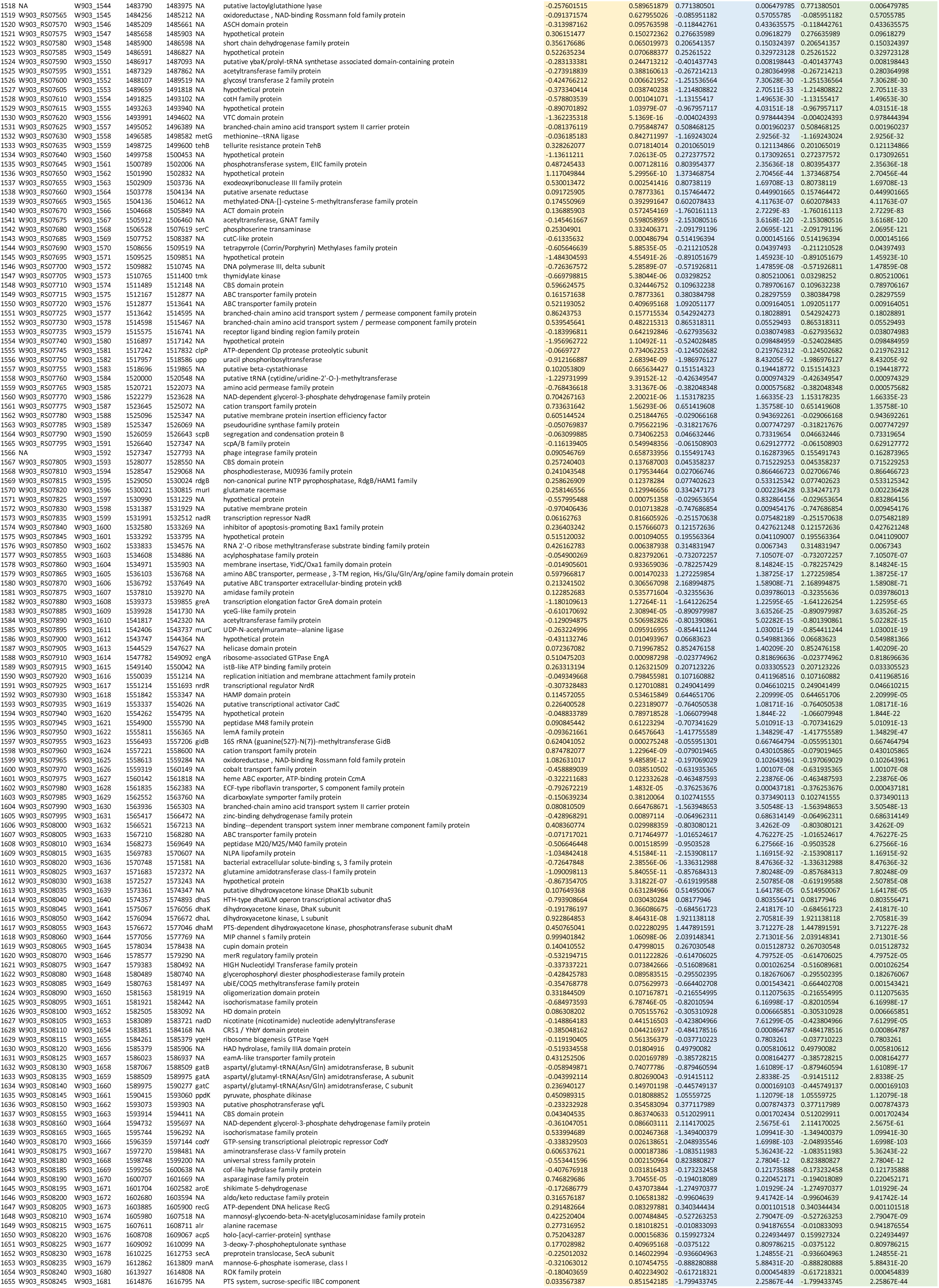
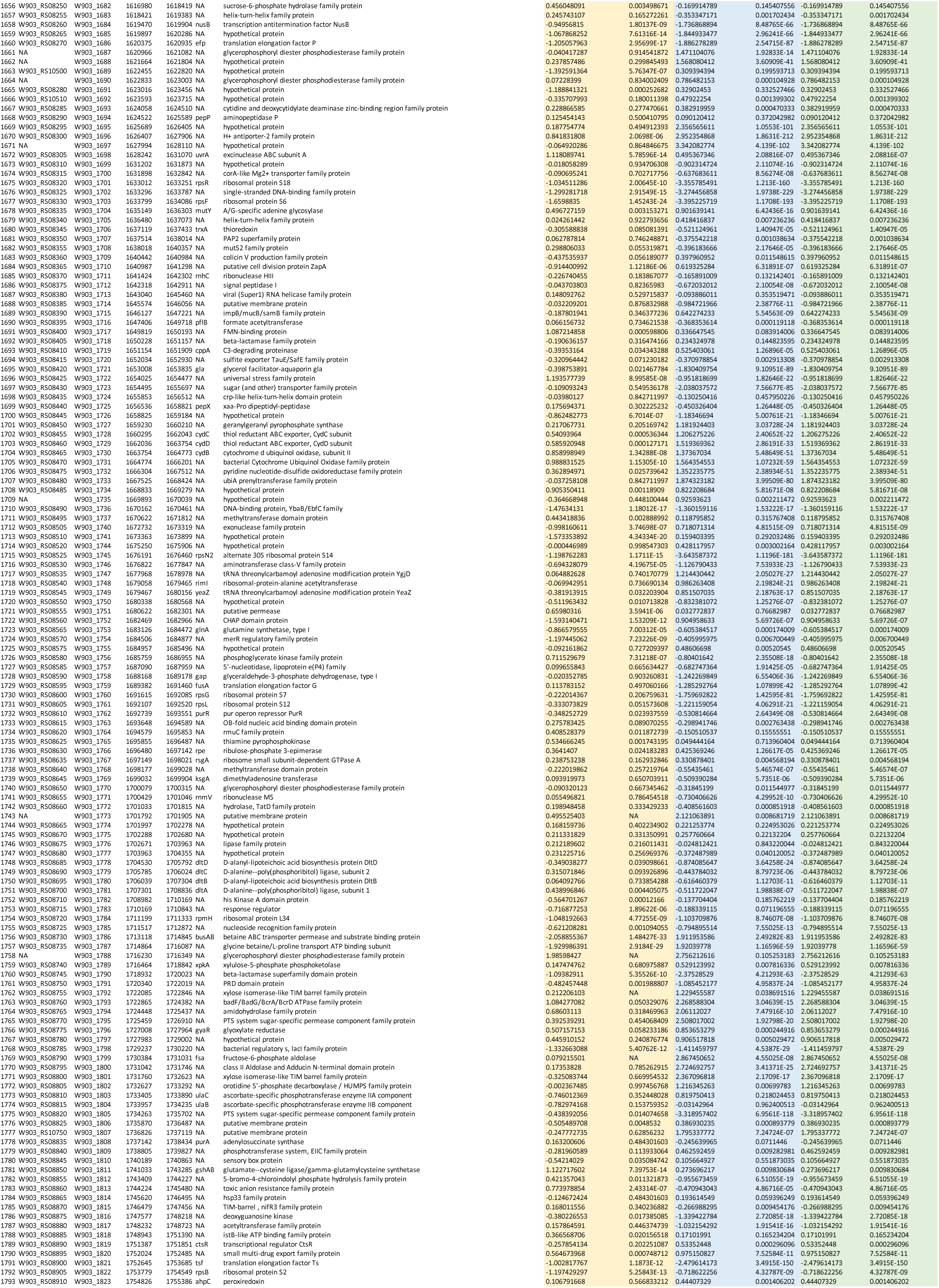
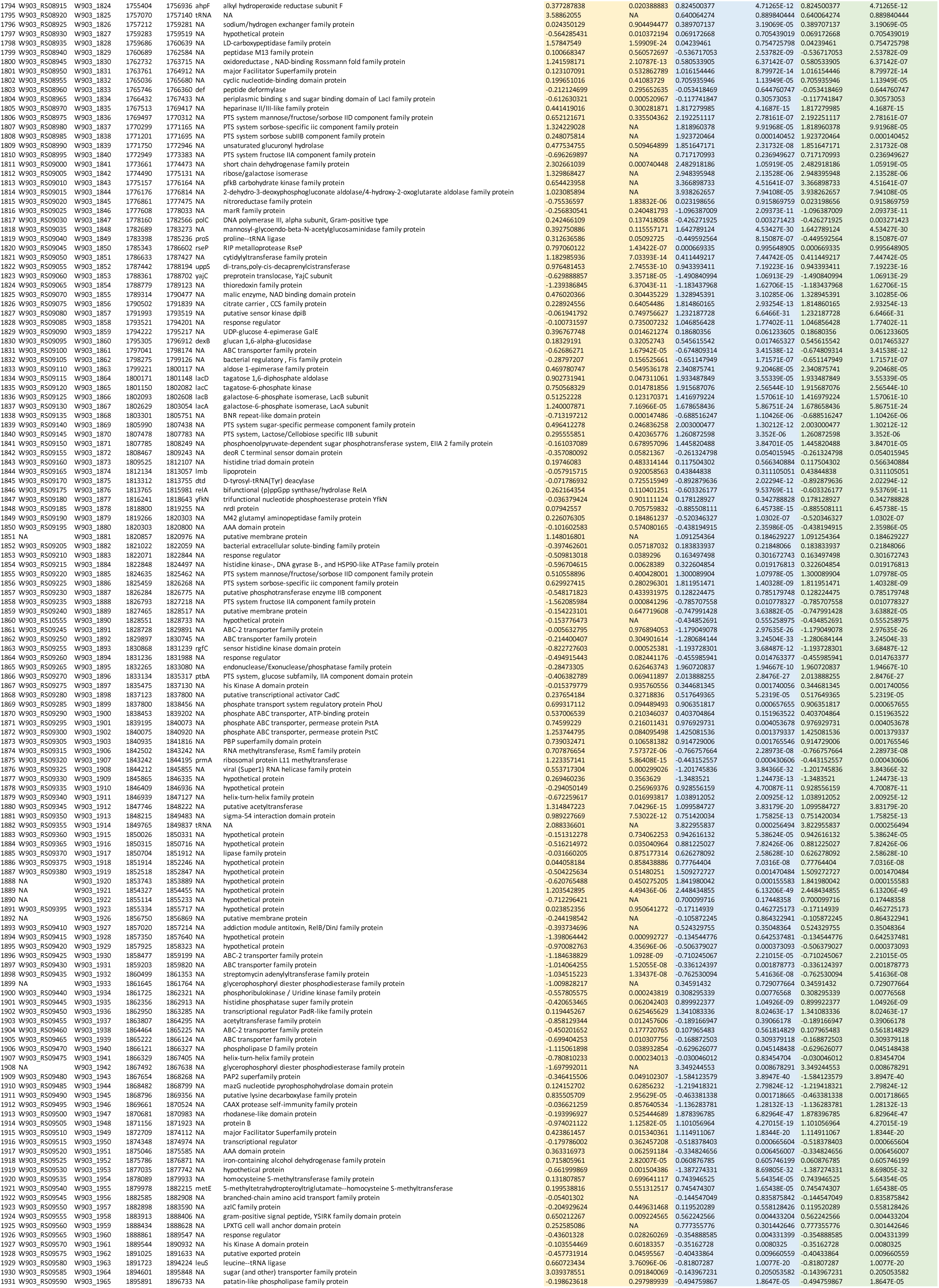
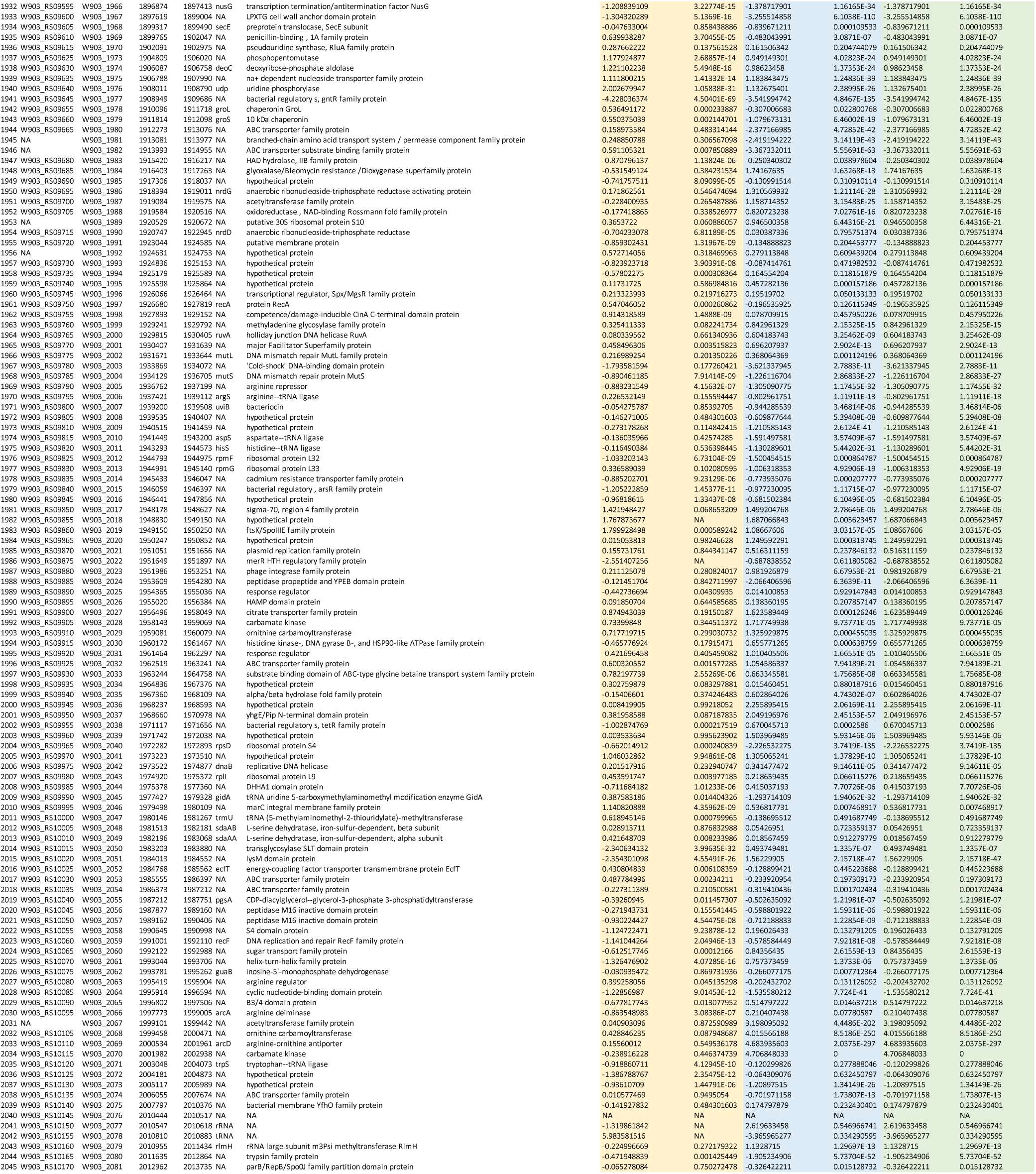

**Supplemental Figure 1:**
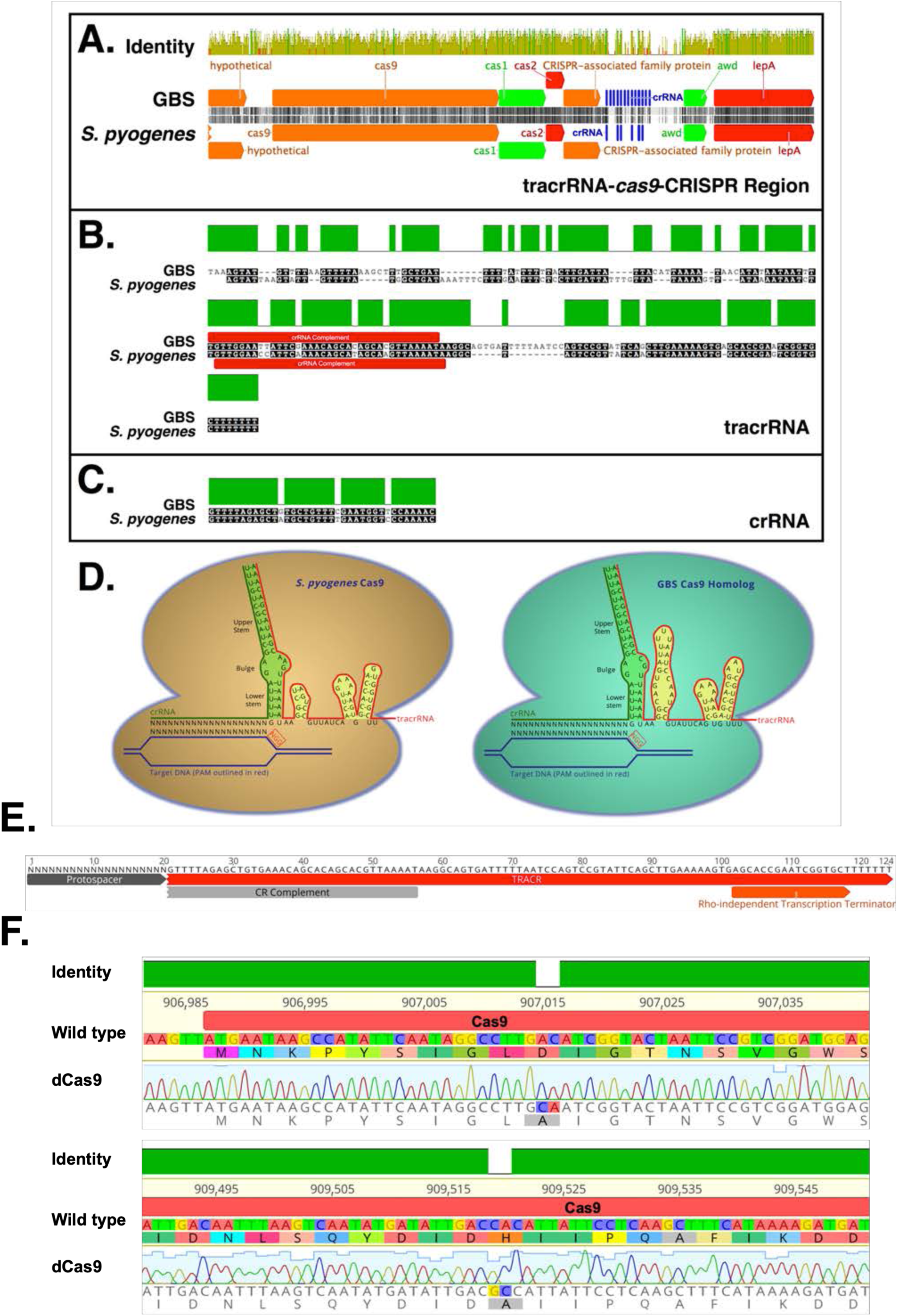
GBS CRISPRi sequence data. Alignments of GBS and *S. pyogenes* genomic DNA regions encoding CRISPR-Cas complex components (**A-C**). Predicted folding of gRNA complexes (**D**) generated using the mfold server (103). Sequence of sgRNA complex used to target candidate genes using GBS CRISPRi system (**E**). Sanger sequence showing the two targeted mutations used to generate dCas9 on the 10/84 chromosome (**F**).

**Supplemental Figure 2:**
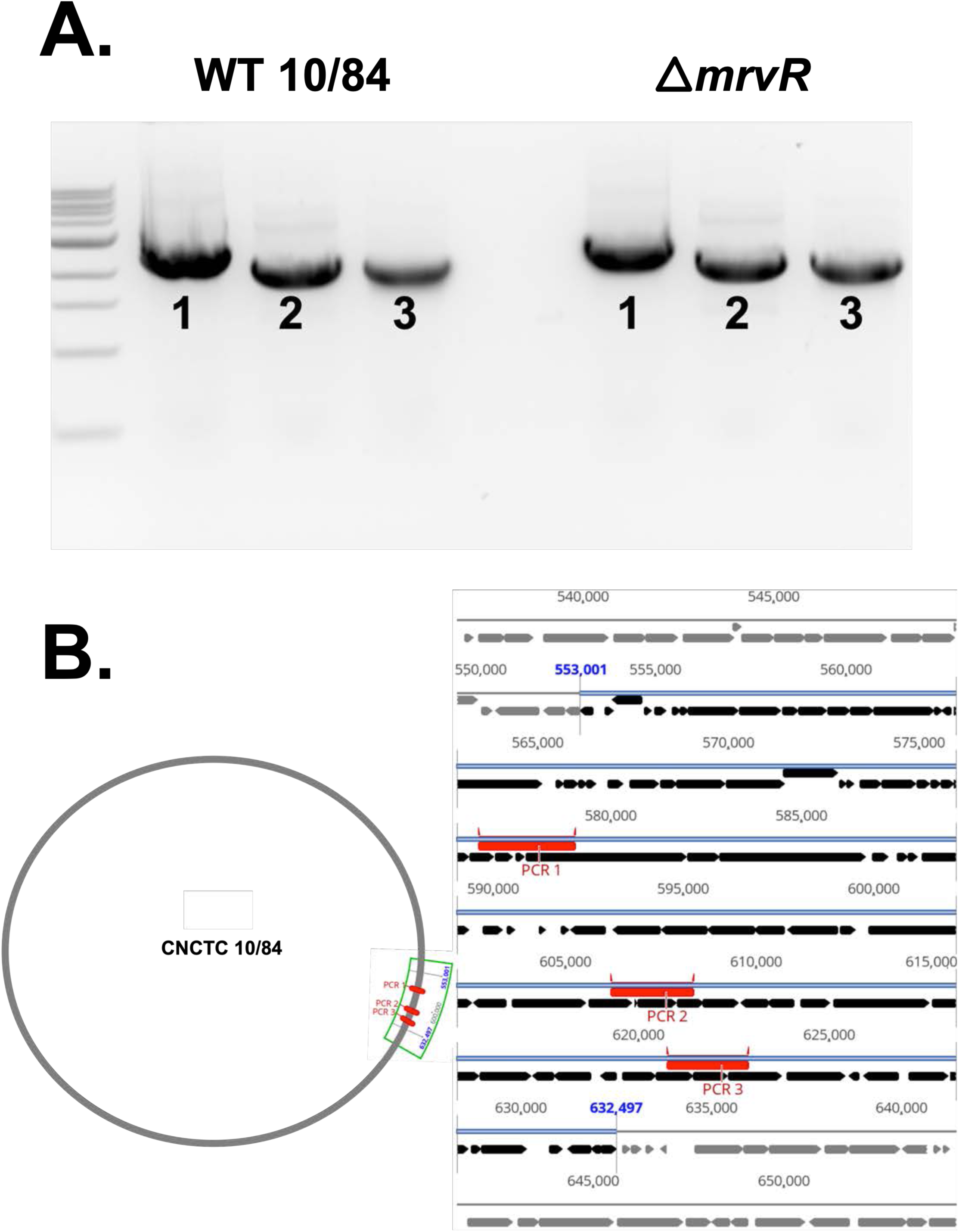
**PCR targeting three regions within the prophage island found to be downregulated in Δ*mrvR*** Three regions within the prophage island spanning gene loci W903_RS03075 through W903_RS03520 were amplified from wild type and *mrvR* knockout GBS genomic DNA. The PCR products from the two strains were the same size when assessed by gel electrophoresis (**A**). The bottom schematic shows the three amplified regions of the chromosome in red (**B**).

**Supplemental Figure 3:**
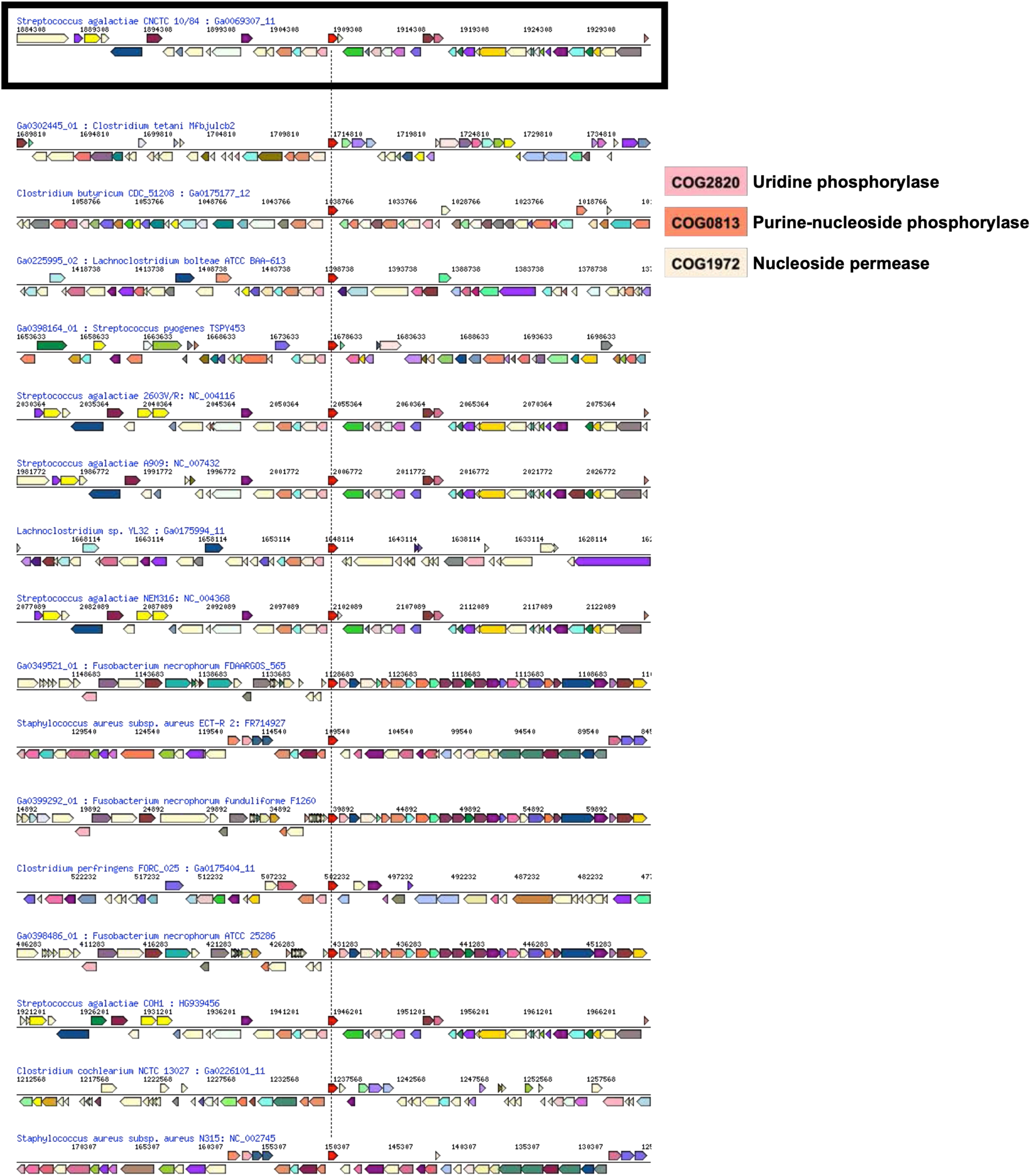
***mrvR* gene neighborhood alignments in other bacteria** Gene synteny plot showing orthologs of the GBS *mrvR* gene in 16 other bacterial species. The plot was generated using the Integrated Microbial Genomes & Microbiomes server on the JGI genome portal (https://img.jgi.doe.gov). Genes are color coded by COG functional prediction. COG categories related to nucleotide metabolism present on the diagram are noted.

**Supplemental Figure 4:**
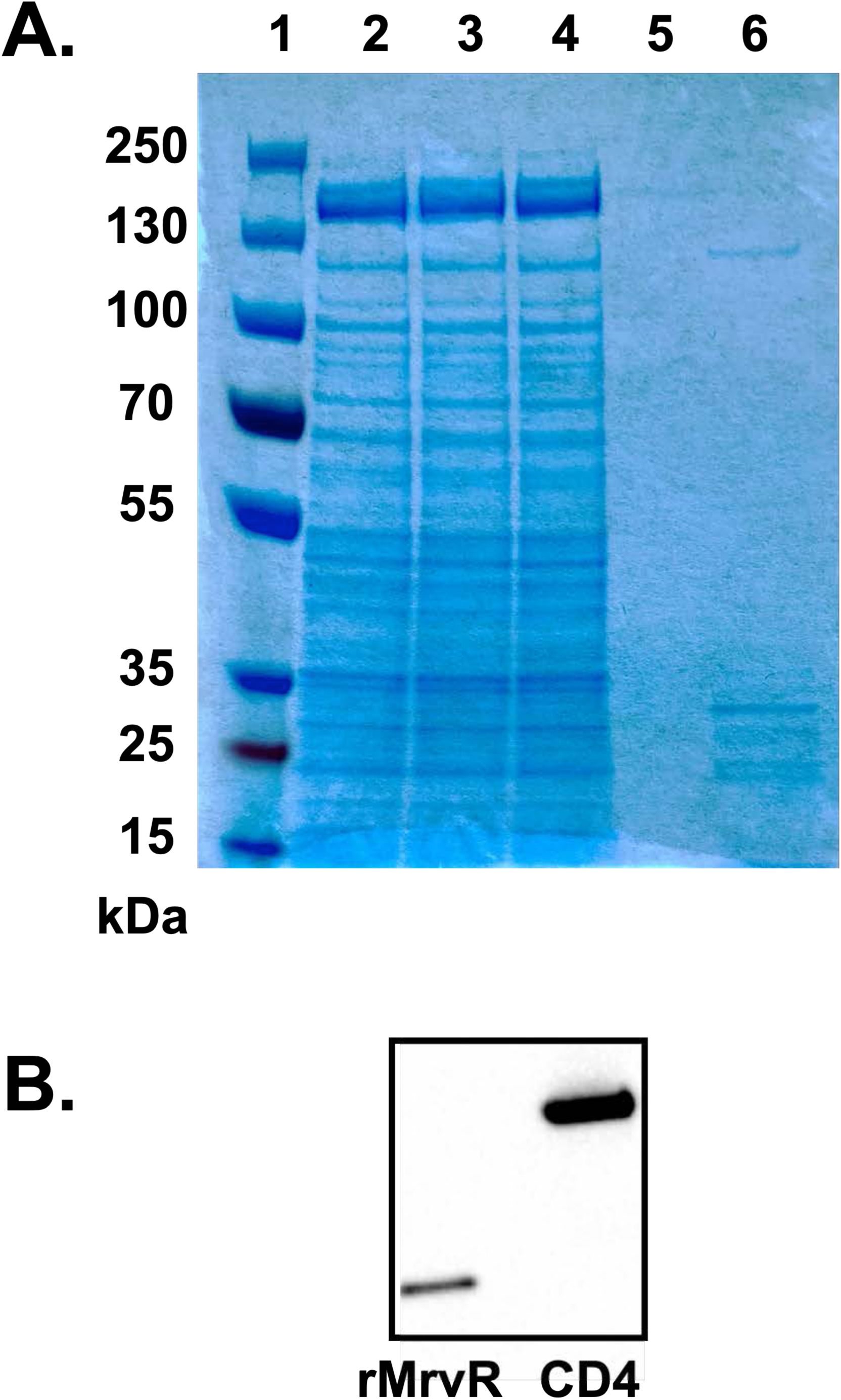
Recombinant MrvR purification. Coomassie stained PAGE gel (**A**) and Western blot (**B**) probed with anti-His tag antibody. Lanes 1-6 in panel A are: Thermo PageRuler Plus prestained protein ladder (cat. # 26620); supernatant from the *Brevibacillus* outgrowth media; supernatant after passage through a 0.2 µm sterilization filter; initial flowthrough from the Takara Capturem his tag purification unit; flowthrough following sequential washes of the purification unit; elution of the purified protein with expected MW=29 kDa.

